# MMORF—FSL’s MultiMOdal Registration Framework

**DOI:** 10.1101/2023.09.26.559484

**Authors:** Frederik J. Lange, Christoph Arthofer, Andreas Bartsch, Gwenaëlle Douaud, Paul McCarthy, Stephen M. Smith, Jesper L. R. Andersson

**Affiliations:** Wellcome Centre for Integrative Neuroimaging, FMRIB, Nuffield Department of Clinical Neurosciences, University of Oxford, United Kingdom; Department of Neuroradiology, University of Heidelberg, Germany

**Keywords:** registration, nonlinear, multimodal, volumetric, FSL

## Abstract

We present MMORF—FSL’s MultiMOdal Registration Framework—a newly released nonlinear image registration tool designed primarily for application to MRI images of the brain. MMORF is capable of simultaneously optimising both displacement and rotational transformations within a single registration framework by leveraging rich information from multiple scalar and tensor modalities. The regularisation employed in MMORF promotes local rigidity in the deformation, and we have previously demonstrated how this effectively controls both shape and size distortion, and leads to more biologically plausible warps. The performance of MMORF is benchmarked against three established nonlinear registration methods—FNIRT, ANTs and DR-TAMAS—across four domains: FreeSurfer label overlap, DTI similarity, task-fMRI cluster mass, and distortion. Results show that MMORF performs as well as or better than all other methods across every domain—both in terms of accuracy and levels of distortion. MMORF is available as part of FSL, and its inputs and outputs are fully compatible with existing workflows. We believe that MMORF will be a valuable tool for the neuroimaging community, regardless of the domain of any downstream analysis, providing state-of-the-art registration performance that integrates into the rich and widely adopted suite of analysis tools in FSL.

## 1. Introduction

In this paper we describe and evaluate FSL’s MultiMOdal Registration Framework (MMORF). MMORF is a nonlinear image registration tool, primarily intended for magnetic resonance imaging (MRI) of the brain.

Biomedical image registration is a core component in most neuroimaging processing pipelines. If we assume that all brains, regardless of appearance, are built using the same anatomical components arranged in the same configuration (*i.e*., they are topologically identical), then we can use image registration to find the set of deformations that map all brains to a common reference space/template brain in a one-to-one manner. In this case, a template may refer to either a group-average brain or an individual subject brain. Although this assumption may not always hold, it is valid enough for registration to be used in both the localisation and quantification of similarities and differences between individuals or population groups—that is, the template allows us to say *where* the differences occur, and the mappings ensure that *what* we are comparing is measured at the same place in all subjects. A poor registration will, therefore, impact both the power to detect and the ability to accurately localise any effects of interest across subjects.

However, the reality is that image registration is an ill-posed problem, and therefore the true, one-to-one mapping can generally never be found. By ill-posed, we mean that there is typically not enough information in an image itself to find a mapping that uniquely maximises some measure of similarity between a subject’s brain and the template. Therefore, registration methods require regularisation to constrain the solution to be unique. Regularisation achieves this by encoding a model of which types of deformations are considered more likely than others. The challenge in designing a registration tool is then to find the best way to combine *image information* and *regularisation* to produce as good an approximation to the true mapping as possible.

MMORF addresses the image information aspect of this challenge by taking a multimodal approach to computing brain similarity. Up until now we have described registration as if there is only one measurement/image of the brain that we can use to find the correct mapping. However, with MRI we are able to acquire a number of different image modalities—all within the same imaging session, and each with different contrast and information content. In MMORF, we have leveraged this complementary information to reduce the degree to which the registration problem is ill-posed, thereby improving our confidence that we are finding an accurate mapping for each subject. MMORF’s registration abilities extend beyond scalar modalities to include diffusion tensor imaging (DTI). When using the full tensor data (rather than scalar, rotationally-invariant, derived features such as fractional anisotropy (FA)), it matches the directional information in the diffusion tensor to guide the alignment of, in particular, white matter.

MMORF’s regularisation method is one of its most unique attributes. By employing a penalty that aims to preserve the original shape and volume of the data as far as is reasonable, it is able to produce deformations that are far more biologically plausible than those generated by conventional techniques. A detailed description of its implementation in MMORF, and a thorough evaluation of the benefits of this form of regularisation, can be found in Lange et al. (2020).

The cost of combining multiple modalities with a complex regularisation model is that the computational requirements of the method increase. We have addressed this from the outset by designing MMORF to use graphics processing units (GPUs) to parallelise its execution. This allows MMORF to execute 1 mm isotropic registrations with reasonable runtimes of between 5 and 45 minutes on modern hardware, depending on the number of modalities used.

Through reducing the degree to which the registration problem is ill-posed (using multimodal data), and producing more realistic deformations in those regions where it is (using biomechanically realistic regularisation), MMORF is capable of state-of-the-art registration accuracy. In the methods section below we detail the most important design decisions made when developing MMORF, so that how it operates can be clearly understood. We then contextualise these decisions with reference to a set of comparable current registration tools. Finally, we validate and benchmark MMORF against these tools in order to demonstrate its performance and utility.

## 2. Methods

As stated in the introduction, the true one-to-one mapping between brain images can never be known, and therefore image registration can only find the “optimal” mapping based on the available information and our prior beliefs about what mappings are more likely than others. Unsurprisingly then, registration is normally formulated as an optimisation problem, requiring a cost function to minimise. MMORF is intended to be used with any number of scalar and tensor images driving the registration, and we therefore choose to minimise a total cost function 𝒞 _*tot*_ that is defined as follows:

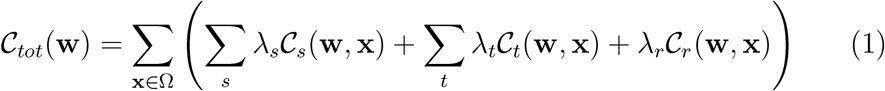

where **w** are the warp parameters being optimised, Ω is the domain over which the warp is defined, *λ*_*_ are cost function weightings, and subscripts *s, t*, and *r* refer to scalar, tensor and regularisation respectively.

Each scalar and tensor cost function is based on an image dissimilarity metric between a pair of images—one belonging to the reference subject (of-ten a template), and one to the moving subject. Although Equation 1 shows that these cost functions are separable, and can therefore be evaluated in-dependently, it is critical that *within* each subject all images are accurately co-registered. If not, then a single warp cannot correctly map all modalities between subjects, and registration accuracy will suffer. We do not attempt to ensure this within MMORF itself. Rather, we rely on running accurate, rigid, *between*-modality, *within*-subject registration using FSL FLIRT (Jenkinson and Smith, 2001; Jenkinson et al., 2002), as well as distortion correction for relevant modalities such as DTI (Andersson et al., 2003; Andersson and Sotiropoulos, 2016), before performing nonlinear registration. However, it is not necessary to resample any of the images following rigid alignment before feeding them into MMORF. Instead, MMORF expects a separate affine transformation matrix to be supplied for each image being registered (in both the moving and reference subject datasets). This affine transformation points to a separate, user-specified, “warp-space image”, whose extents define the domain over which MMORF will estimate and output its nonlinear deformation. This eliminates the need for multiple resampling of the data, and requires no matching of the image resolution or dimensions of any of the images being registered.

With this information in hand, the following sections elaborate on some of the key decisions that went into MMORF’s design, and the role they play in its performance.

### 2.1. Optimisation Strategy

Nonlinear image registration tools generally use one of two iterative optimisation approaches—first or second order. First order methods consider only the gradient of the cost function, with respect to each optimisable parameter, when picking a parameter update step. Second order methods extend this by also considering how long the derivative is valid for and the interaction between parameters. First order algorithms tend to have update steps that can be calculated more quickly, but second order algorithms tend to require fewer update steps to reach convergence. In our experience, the trade-off tends to favour second order approaches for methods such as MMORF.

We have, therefore, implemented two variants of the (second-order) Gauss-Newton (GN) optimisation strategy—which is itself a variation on Newton’s method.

Newton’s method is an iterative optimisation algorithm that uses a quadratic Taylor approximation of the cost function 𝒞 around the current set of parameters **w** (Nocedal and Wright, 2006, Ch.10, pp 254). That is:

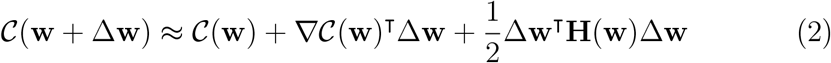

where ∇𝒞 and **H** are the gradient and the Hessian of the cost function, respectively. The update Δ**w** that minimises this approximation to the cost function (when the cost function is convex) is then:

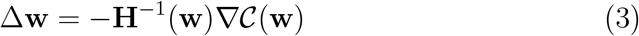

For cost functions that can be written in the form:

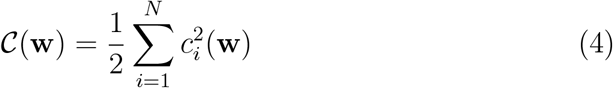

The gradient is then:

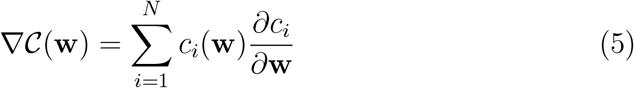

And the Hessian is:

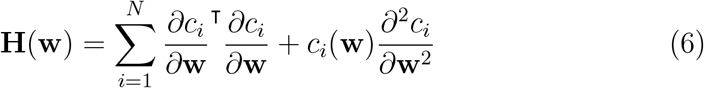

Gauss-Newton approximates **H** by dropping the second, mixed partial derivative, term in Equation 6. Close to the optimum set of **w**, the Newton and Gauss-Newton Hessians tend towards equivalence. Further from the optimum, the Gauss-Newton Hessian has the benefit of being symmetric positive definite, and therefore the approximation of the cost function is always convex. This ensures that each step is always in a direction that would improve the original cost function.

However, the Gauss-Newton step can still lead to an increase in the cost function if it oversteps. This can be addressed by the Levenberg-Marquardt (LM) extension (Nocedal and Wright, 2006, Ch.10, pp258), which replaces the Gauss-Newton Hessian (**H**_*GN*_) with:

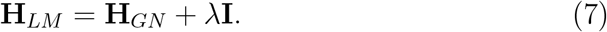

For small values of *λ*, the update Δ**w** behaves much like a Gauss-Newton update, and for large values of *λ* it behaves more like gradient descent. After choosing a sensible starting value for *λ* (e.g., 0.001), a typical Levenberg-Marquardt iteration then proceeds as follows:

- Calculate a candidate parameter update Δ**w** using Equation 3.
- If 𝒞 (**w** + Δ**w**) *<* 𝒞 (**w**) accept the update and set *λ* = *λ ÷* 10.
- If 𝒞 (**w**+Δ**w**) *>* 𝒞 (**w**) set *λ* = *λ*×10 and recompute Δ**w** and 𝒞 (**w**+Δ**w**), repeating until 𝒞 (**w** + Δ**w**) *<* C(**w**) and then accept the update.

The Majorise-Minimisation (MM) algorithm (Hunter and Lange, 2004) is a method for cost function minimisation that can be used when storing the full Gauss-Newton Hessian is infeasible (*e.g*., due to memory constraints, which prove to be important in this application). In essence, what this algorithm states is that if *z*(**w**) ≥ *y*(**w**), ∀**w**, with both *y* and *z* convex and touching, then **w**_*k*+1_ = arg min *z*(**w**) will either reduce *y* or leave it unchanged.

Since the quadratic Taylor approximation of 𝒞 (**w**) using **H**_*GN*_ in Equation 2 is convex, it may serve as *y*. Chun and Fessler (2018) showed that because the diagonal elements of **H**_*GN*_ are all positive:

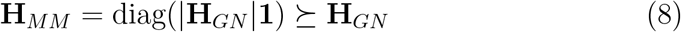

where |**H**_*GN*_ | is the matrix of the absolute values of **H**_*GN*_ and **1** is column vector of ones. The ⪰ symbol means that **H**_*MM*_ is at least as positive definite as **H**_*GN*_ . Therefore, substituting **H**_*MM*_ for **H**_*GN*_ in the quadratic Taylor approximation of 𝒞 (**w**) majorises *y*, and may serve as *z*. In other words simply, replacing **H**_*GN*_ with **H**_*MM*_ in the LM algorithm and solving as usual satisfies the MM algorithm requirements.

The major advantage of using **H**_*MM*_ is that it is only non-zero along its main diagonal, and therefore requires far less memory to store than **H**_*GN*_ . This is particularly important when the number of parameters in **w** is large, as it may not be possible to store **H**_*GN*_ .

The reason for including both the Levenberg-Marquardt and Majorise-Minimisation modifications of Gauss-Newton, is that MMORF both calculates the Hessian and solves for the update step on GPU hardware. GPUs typically have a limited amount of RAM, and, therefore, storing the full Gauss-Newton Hessian becomes impossible for warps beyond a certain resolution. At this point, MMORF switches to Majorise-Minimisation, which requires on the order of 1000 times less memory. In practice, Majorise-Minimisation requires more steps to converge, but requires no other changes to the registration algorithm, making it an appealing option.

### 2.2. Transformation Model

MMORF employs a transformation of the small deformation framework variety (Bajcsy and Kovačič, 1989; Miller et al., 1997)—that is, it defines the deformation as a displacement field, rather than a velocity field as per the large deformation framework (Bro-Nielsen and Gramkow, 1996; Miller et al., 1997). A major difference between these two frameworks is their relationship to diffeomorphism.

Diffeomorphism is a desirable property in image registration. Diffeomorphic transformations are smooth, one-to-one, and onto, and are therefore guaranteed to induce neither folding nor tearing when applied to an image. The large deformation framework, and in particular the large deformation diffeomorphic metric mapping (LDDMM) (Beg et al., 2005) family of tools, have the advantage that their transformation model can be made inherently diffeomorphic—i.e., they are, in principle, diffeomorphic by construction.

However, despite not being inherently diffeomorphic, the small deformation framework can be made diffeomorphic by employing a regularisation penalty that enforces diffeomorphism (Rohlfing et al., 2003). This has the benefit that warp-induced distortions, such as changes in shape and volume, can be calculated (and therefore controlled) directly from the model parameters themselves. In contrast, large deformation models require the vector field that they parametrise to first be integrated over time and converted into a displacement field before being able to calculate such measures. Therefore, explicitly *controlling* the amount of distortion is harder in the large deformation framework, despite its guarantee of diffeomorphism. This is important in our case, as the advantage of MMORF’s regularisation (described in Section 2.4) is that it nonlinearly penalises the stretching and compressing effects of the warp directly.

MMORF’s transformation is parametrised using cubic B-splines (Unser et al., 1993a,b). This imposes an inherent smoothness to the deformations, as cubic B-splines, and therefore the warps, have C2 continuity. The warp field is then also well defined across all of the image (*i.e*., not just at voxel centres), eliminating any need to match image subsampling or resolution either before or during registration. Despite their smoothness, B-splines still have compact support (*i.e*., the effect of a particular spline is exactly zero at a fixed distance from the spline centre), and therefore interaction effects between splines are fixed and finite. This has important implications for second order optimisation (such as the Gauss-Newton method employed by MMORF), since it means that the Hessian matrix (which encodes the interaction between optimisable parameters) is sparse and predictably patterned. Furthermore, the spatial derivatives of B-splines are smooth and have a closed form solution. Since these derivatives are required to calculate gradients and Hessians for the optimisation of warps in MMORF, this is computationally beneficial.

We now provide a more explicit description of how we have formulated our transformation model, and how this interacts with calculation of the Hessian during optimisation (the most computationally intensive part of the registration algorithm). A set of B-spline basis functions can be used to transform a set of sample coordinates in a reference image *f* to their corresponding coordinates in a moving image *g* as follows:

*f* = Reference image

*g* = Moving image

*N* = Number of sampled voxels in *f*

**x***/***y***/***z** = *x, y* and *z* coordinates of samples in *f*

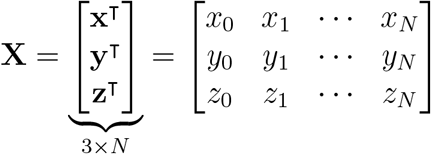

*M* = Number of splines per warp direction

**w**_*x*_*/***w**_*x*_*/***w**_*z*_ = *x, y* and *z* direction warp parameters

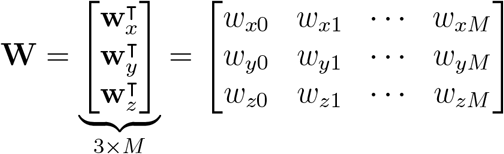

**b**_*m*_ = Vectorised *m*^th^ B-spline at sample positions **X**

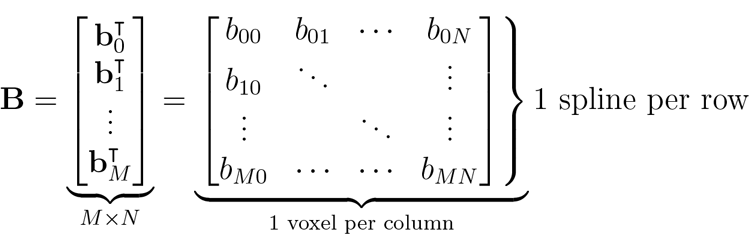

*ϕ* (**X, W**) = Transformed sample coordinates

= **X** + **WB**

Note that here we assume that the warp space and the reference image *f* are the same, and that *f* and *g* are already affinely registered to each other. We use the same configuration of B-splines to define the warps in all 3 directions (*i.e*., the number, order and spatial extent of splines defining the displacements in each direction (*x, y* and *z*) is the same, and they are located identically in space). *M* is chosen to be the set of splines whose spatial support, wholly or partially, overlaps with the domain over which *f* is defined. The warp parameters **w**_*x/y/z*_ are then the coefficients of each Bspline, and each parameter only affects the displacement in a single direction.

The compact support of B-splines means that **B** is very sparse. Additionally, (disregarding edge cases) each row of **B** is simply a shifted version of any other row. We therefore never store **B** explicitly, and instead compute **WB** using convolution.

Another benefit of the sparsity in **B** is that it induces sparsity in the Hessian of cost functions based on this transformation. For a mean squared error cost function, each element of the Hessian can be calculated as:

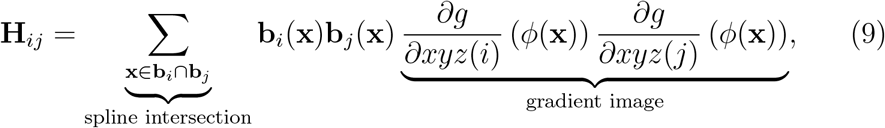

where 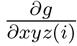 means “differentiate with respect to the direction in which Bspline *i* causes displacement to occur”. As each entry in the Hessian represents the interaction between two of the B-spline basis functions, only those combinations of splines that overlap in their spatial support will ever produce non-zero entries. The number of non-zero entries per row of the Hessian is at most 3 ×7^3^ = 1026 for 3D images and cubic B-splines. Considering that the number of parameters being optimised over can easily exceed 10^6^, this would lead to a matrix that is at least 99.9% sparse. The redundancy increases as the warp resolution increases (the knot-spacing is reduced), which means that the more parameters one attempts to estimate, the sparser the Hessian becomes. Therefore, the memory requirement for the Hessian scales much better with resolution than might initially be feared.

### 2.3. Image Cost Functions

As stated earlier, MMORF optimises the cost function defined in Equation 4, which is the sum over individual cost functions for each pair of images. In this section we describe the choice of cost function for scalar and tensor images, as well as how MMORF implements cost function masking/weighting.

#### 2.3.1. Scalar

Scalar image dissimilarity is calculated using the mean squared error (MSE) across the image. That is, for two scalar images *f* and *g*:

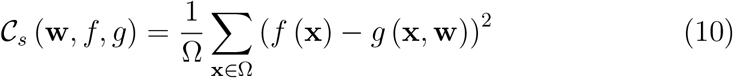

MSE is ideally suited to the GN family of optimisation methods, which have an implicit assumption that the cost function being optimised is some form of squared difference.

Robust mean intensity estimation is used to scale each image separately in order to account for linear scaling intensity differences between image pairs. When using MSE cost functions, spatial intensity inhomogeneities (*e.g*., due to transmit and receive bias fields in MRI (Vovk et al., 2007; Andersson et al., 2019)) can be mistakenly interpreted as image misalignment, leading to unnecessary (and incorrect) image warping. In MMORF we therefore provide the option to explicitly model such inhomogeneities as a smoothly varying multiplicative field acting on the reference image. This can be enabled or disabled for each image pair independently. The bias field is parametrised using cubic B-splines, and the resolution (knot-spacing) and smoothness (bending energy (Bookstein, 1997)) are set on a per image pair basis.

#### 2.3.2. Tensor

Tensor image dissimilarity is calculated using the mean squared Frobenius norm (MSFN) across the image. This is exactly equivalent to summing the MSE for each of the 9 elements of the diffusion tensor, and therefore fits our GN optimisation strategy. That is, for two tensor images **F** and **G**:

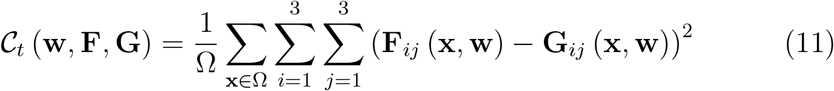

Note that, in contrast to the scalar case, the values in the reference image **F** are also a function of the warp parameters **w**. This is because MMORF applies the *rotational* effect of the warp only to the reference image, as well as the usual *displacement* effect only on the moving image.

Rotation of tensors uses the finite strain (FS) method (Alexander et al., 2001). The Jacobian matrix of the warp at each position **J**(**x, w**) represents the first order linear approximation of the deformation at that point, and from it the local rotational effect of the warp **R**(**x, w**) can be calculated using:

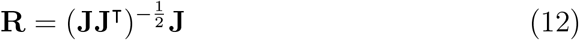

We include the effects of both displacement and rotation when calculating the gradient and the Hessian of the tensor cost function, using the derivation of Yeo et al. (2009) for the closed-form of the derivative due to rotation.

Since the diffusion tensor model is quantitative, no intensity rescaling is performed. This implicitly assumes that the tensors are always represented with the same units. Similarly, fitting of the diffusion tensor compensates for any intensity inhomogeneity present in the raw diffusion-weighted images, and therefore no bias field estimation is required .

MMORF assumes that diffusion tensors are stored in FSL dtifit1 format—*i.e*., a 4D volume where the 4^th^ dimension contains the 6 upper-diagonal elements vectorised row-wise, and the x-direction is defined in radiological convention (R-L).

#### 2.3.3. Masking

In certain instances it may be beneficial to focus a registration algorithm on a particular part of an image or, alternatively, for it to ignore a certain region. To this end, we have implemented masking/weighting within MMORF.

Masks can be supplied for all reference and moving images independently, and it is required that these masks be in the same space as their corresponding images. The masks are treated as containing voxelwise multiplicative factors that are applied to the cost function during optimisation. Setting any region of a mask to zero will cause the algorithm to ignore the impact of that region’s similarity between reference and moving images, and the deformation in that region will be determined purely by the regularisation. However, masks do not have to be binary, and so a “soft” mask can be used to favour the alignment of one region (*e.g*., the brain over the rest of the head), without ignoring it completely.

In MMORF, masks can be enabled or disabled for each image at every iteration independently, allowing for maximum flexibility. We have, at times, found it useful to have a mask be applied only during higher resolution iterations of the registration.

### 2.4. Regularisation

Since image registration is an under-constrained problem, regularisation is essential to ensure that the resulting warp fields are biologically plausible.

A plausible warp should, at the very least, be diffeomorphic. Diffeomorphic warps are both one-to-one and onto—that is, each point in the reference space maps to a unique point in the moving space, and each point in the moving space is reachable from the reference space. They require the mapping to be continuous, smooth (*i.e*., have a continuous derivative), and invertible (*i.e*., have a finite, positive Jacobian determinant everywhere). B-spline parametrised warps are, by definition, smooth and continuous. Therefore, provided the Jacobian determinant remains *>* 0 everywhere, the warp is diffeomorphic.

Diffeomorphism is, however, only one desirable trait in a warp. It guarantees that an image is never torn or folded, but that is all. Typically, penalising the variation of the Jacobian determinant from a value of 1 is used to encourage or ensure (depending on the choice of penalty) that warps remain diffeomorphic in the small deformation framework. However, penalising the Jacobian determinant directly only controls volumetric distortion (*i.e*., changes in an object’s size), and does not in any way control shape distortion (*i.e*., changes in an object’s shape). The singular values of the local Jacobian represent stretches/compressions along three orthogonal directions. Therefore, the difference between them indicates the degree of shape distortion. Because the Jacobian determinant is the product of the singular values of the Jacobian, one can control changes in both size and shape by controlling only the singular values. Penalising deviations of the singular values from 1, and ensuring that none become negative, leads to diffeomorphic warps with desirably little distortion in both volume and shape.

In MMORF, the specific penalty used is the mean (across the image) of the sum of the squared logarithms of the singular values at each voxel, as shown in Equation 13 where *s*_*i*_ is the *i*^th^ singular value of the local Jacobian matrix **J**. This is an adaptation of the penalty first proposed by Ashburner et al. (1999, 2000). Its implementation in MMORF is described (and evaluated) in detail in Lange et al. (2020), where we demonstrate the positive effect that this form of regularisation has on the biological plausibility of the warps MMORF produces.

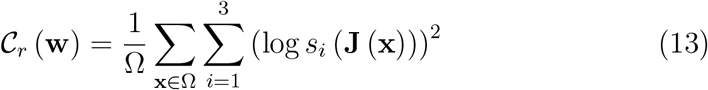

By taking the squared logarithm, this penalty tends to infinity as any singular value tends to either zero or infinity. Additionally, this does not penalize any transformations that are locally rigid (*i.e*., regions that are only translated and/or rotated). Therefore, MMORF regularisation enforces diffeomorphism and encourages local rigidity thereby controlling both volumetric and shape distortions. The highly nonlinear form of the regularisation also allows the weighting to be set such that large deformations are allowed when necessary, while still ensuring diffeomorphism, thereby overcoming one of the perceived limitations of the small deformation framework.

### 2.5. Inverse Consistency

An inverse consistent registration algorithm will produce the inverse of the original warp when the reference and moving images swap roles (Christensen, 1999). Since we often consider the choice of reference and moving image to be arbitrary (*e.g*., such as when registering two individuals to each other), this is a desirable property to have.

Some tools, such as ANTs (Avants et al., 2008), are inverse consistent by design (excluding the affine initialisation). This is achieved by registering both moving and reference images to a mid-space. However, this essentially means running two registrations per pair of subjects, which takes double the time to compute.

In MMORF we have taken a different approach where, rather than enforcing a symmetric warp, we symmetrise the cost function being minimised instead. This is achieved by multiplying each cost function (both image similarity and regularisation) by a weighting term of 1 + |**J**| . Since the cost functions are evaluated on a regularly sampled grid in the space of the reference image, the intuition for this weighting is that it always accounts for the total volume in both images to which that value of the cost function applies. In the continuous case, this can been shown to exactly symmetrise the cost function (Tagare et al., 2009). However, since we are dealing with discretely sampled data, this correction is only approximate in MMORF. Nevertheless, this is a better solution than leaving symmetry unaccounted for, and has the computational benefit that only one warp field need be calculated during registration.

### 2.6. Multiresolution Pyramid

MMORF employs a coarse-to-fine multiresolution optimisation strategy (Bajcsy and Kovačič, 1989; Szeliski and Coughlan, 1997). This has been shown to be beneficial in avoiding local minima during optimisation, as well as accelerating convergence, across a wide variety of nonlinear registration tools (Zhang et al., 2006; Andersson et al., 2007; Ashburner, 2007; Avants et al., 2008; Modat et al., 2010). In principle, such approaches try to match low-frequency image information first, followed by progressively higher frequency information at each subsequent level. Traditionally, users are required to specify a downsampling factor and amount of smoothing for each level of the pyramid. The lower-resolution warp is then defined on the coarser, downsampled reference image grid. A potential pitfall of this approach is that if insufficient smoothing is applied to the image, then the process of downsampling introduces aliases in the information being aligned due to violation of the Nyquist criterion. MMORF overcomes this by defining the pyramid according to warp resolution (knot spacing) and image smoothing only. An acceptable level of subsampling is then automatically determined by treating the Gaussian smoothing kernel as a low-pass frequency filter. Therefore, the user can iteratively optimise the registration of different frequency content within the image through applying decreasing amounts of smoothing while keeping the warp resolution the same, without any problems with aliasing. In practice, we restrict the downsampled resolution to be at least as high as, and at most four times higher than, the warp resolution at each level of the pyramid.

### 2.7. Other Implementation Considerations

MMORF is written in C++ and makes extensive use of GPU parallelisation using Nvidia’s CUDA framework (NVIDIA, 2019). Without the use of GPU parallelisation, certain aspects of the registration would be too computationally burdensome to allow MMORF registrations to complete within a reasonable runtime. With GPU acceleration, a typical 1 mm isotropic registration with MMORF takes ≈ 10 min for a single scalar, unimodal image pair and ≈ 45 minutes for a single scalar, single tensor, multimodal image pair. However, this reliance on CUDA means that MMORF is only supported on Linux devices with Nvidia GPUs. MMORF uses a mixed computation model, with only the most time-consuming components of the registration algorithm being ported to the GPU. These include

- Image interpolation
- Cost, gradient and Hessian calculations
- Solving the system of linear equations for update steps

Of these, the Hessian calculation is by far the most computationally complex, with those of the regularisation and the rotational component of the DTI cost functions being particularly burdensome. For these calculations, GPU acceleration is on the order of 20-40x (depending on warp resolution, image dimensions and image subsampling (Lange et al., 2020, Supplementary Material: GPU Considerations and Code Profiling)).

MMORF has been designed from the outset for integration into the FSL suite of neuroimaging tools (Jenkinson et al., 2012). As such, it expects inputs in FSL convention. Specifically, affine matrices between input images and the warp space should be in FLIRT format and DTI images should be in dtifit format. MMORF output warp fields follow existing FSL conventions and can therefore serve as drop-in replacements for FSL commands such as applywarp.

## 3. Theoretical Differences Between Methods

With an understanding of the design considerations that went into MMORF, we will now briefly describe how some of these choices compare to three existing registration tools, namely: FNIRT (Andersson et al., 2007), ANTs Avants et al. (2008) and DR-TAMAS (Irfanoglu et al., 2016). These three tools will then be used to validate and evaluate the performance of MMORF in Section 4.

FNIRT was chosen as it is the predecessor to MMORF, and the current nonlinear registration tool in FSL. The largest differences between these methods are MMORF’s multimodal capabilities, regularisation, inverse-consistency and GPU parallelisation. In terms of similarities, they share the same transformation and bias field models, and very similar optimisation strategies at low resolutions. At higher resolutions, FNIRT switches to a Scaled Conjugate Gradient algorithm (Møller, 1993), in contrast to MMORF’s MM approach. Finally, FNIRT performs a simultaneous optimisation of both warp and bias fields, whereas these are optimised in a greedy, interleaved manner by MMORF. This is because simultaneous optimisation results in a Hessian without the regular diagonal structure, on which MMORF relies for efficient GPU parallelisation. In practice, despite some similarities in design choices, they perform very differently—even when MMORF is run, like FNIRT, in a unimodal configuration. The resulting warps have a very different character, which we largely attribute to the superior regularisation metric employed in MMORF. MMORF’s inputs and output files are fully compatible with those of FNIRT, and it can therefore serve as a drop-in replacement in FSL analysis pipelines.

ANTs was chosen due to its consistently high performance, including in dedicated registration comparisons (Klein et al., 2009). It has become a *de facto* standard for nonlinear registration in much of medical imaging, and serves as a benchmark against which to compare MMORF’s performance. ANTs is a purely scalar registration method, although it can be applied to multiple scalar input modalities simultaneously. ANTs uses a symmetric, greedy approximation of large deformation diffeomorphic metric mapping (LDDMM) known as symmetric normalisation (SyN). At each iteration, an update step is composed with the current warp until convergence is reached.. If each update step is diffeomorphic, then the composition of all updates is also diffeomorphic (apart from arithmetic floating point errors). Symmetry is achieved by registering both reference and moving images to a mid-space at each iteration. There are a number of similarity metrics implemented, but we will limit ourselves to locally normalised cross-correlation (LNCC), as this has proven to perform best in previous studies. ANTs regularisation consists of simple Gaussian smoothing, which may be applied independently to both update steps and to the final deformation field.

DR-TAMAS was chosen as it is currently the only other tool, to our knowledge, which can simultaneously register both tensor and scalar modalities in a single framework. It has also proven to match or exceed the best performing diffusion registration tools currently available. As in MMORF, finite-strain reorientation of the tensors is taken into consideration during each update step (*i.e*., it contributes to the gradient of the cost function, and is not simply applied after each update step). In contrast to MMORF, the DTI dissimilarity is divided into two separate cost functions—the trace similarity (a rotationally invariant scalar), and the deviatoric tensor similarity (sensitive to relative tensor orientation). For scalar inputs, DR-TAMAS uses an LNCC cost function. The transformation model and optimisation strategy are the same as ANTs, and therefore DR-TAMAS can be considered as a truly multimodal variant of ANTs. In many ways, then, DR-TAMAS is the natural alternative to MMORF.

There are clearly many other nonlinear registration tools against which we could have compared, but we believe that these choices allow us to effectively benchmark the relative performance of MMORF against those tools that are most likely to be considered as an alternative by users.

## 4. Validation

Benchmarking a registration tool can be difficult to do well. Measures of accuracy are often biased towards metrics based on the modalities that drove the registration (Irfanoglu et al., 2016), and evaluations hence risk some degree of circularity. Methods often perform best when evaluated using metrics based on the same data that drove the registration (Irfanoglu et al., 2016), which risks introducing a degree of circularity in these types of evaluations. We have, therefore, endeavoured to perform a holistic evaluation of registration performance by including a range of structural-, diffusion-, functional- and morphometry-derived metrics. Structural and diffusion metrics are the most circular, since these are the modalities that drive the registration (either individually or jointly) in all methods—however, they may highlight both the value and the pitfalls of using the data you wish to analyse to drive alignment. Functional metrics are based on a fully held-out modality (that is not seen by any of the registration tools) and therefore serves as the best proxy of true consistency of anatomical alignment across individuals—since overfitting to a driving-modality (*e.g*., unrealistically deforming the brain to make structural images appear very similar) is likely to have a negative effect on functional alignment (Coalson et al., 2018; Robinson et al., 2018). Morphometry metrics (*e.g*., measures of distortion that represent how aggressively the images are deformed) are essential to contextualise and interpret the accuracy/similarity metrics—that is, it is important to know how aggressively a tool needs to deform an image to achieve a certain degree of registration accuracy, with less distortion being preferred.

We chose the Human Connectome Project (HCP) young adult 1200 release (100 unrelated subjects subset) dataset as the basis for our testing (Van Essen et al., 2012, 2013; Glasser et al., 2013). The HCP dataset contains high quality T1w (0.7 mm isotropic), diffusion (1.25 mm isotropic) and task-fMRI (2.0 mm isotropic) scans. This allows both unimodal T1w and multimodal T1w + DTI registration to be conducted with the same dataset, as well as evaluating registration metrics based on all three modalities. The minimally preprocessed HCP data were used as far as possible (Glasser et al., 2013), which includes motion and distortion correction with FSL topup and eddy (Andersson et al., 2003; Andersson and Sotiropoulos, 2016), and coregistration of the diffusion data to T1w space. The task-fMRI data were, however, reprocessed in subject T1w space with no smoothing (rather than in MNI-152 space with 2.0 mm isotropic smoothing). The diffusion tensor model was fit to the b=1000 s*/*mm^2^ shell only using FSL dtifit.

The Oxford-MultiModal-1 (OMM-1)^2^ template was used as the reference space for all registrations (Arthofer et al., 2021, 2022). This template was constructed from 240 UK Biobank (Miller et al., 2016) subjects, and has both T1w and DTI volumes that are intrinsically co-registered. The OMM-1 therefore provides a common space in which to compare methods that either use T1w images only, or a combination of T1w and DTI images to drive the registration.

We calculate two structural accuracy metrics based on pairwise label similarity. The first is a measure of overlap (specifically, the Jaccard coefficient) and the second a measure of distance (specifically, the Hausdorff distance) of automatically generated cortical and subcortical labels for each pair of subjects following resampling to template space (see Section 4.2 for details). The Jaccard coefficient for two regions *A* and *B* is defined as:

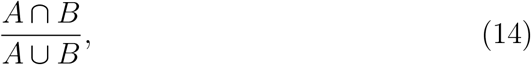

and ranges from 0 for no overlap to 1 for perfect overlap. The Hausdorff distance is the maximum distance between the surfaces of the regions being compared. The labels are derived from the T1w images, and therefore we expect them to favour scalar registration methods that can match the T1w images without being penalised if that reduces DTI similarity. We have avoided any simple intensity-based or tissue-type overlap metrics, as these are known to correlate poorly with true anatomical consistency (Rohlfing, 2012).

We then calculate three diffusion accuracy metrics that compare the similarity of the template DTI modality with each subject’s DTI data after resampling to template space. Each subject is compared voxelwise to the template, and the average across all voxels within a mask is taken as the overall metric for that subject. Overall tensor similarity (OVL, Equation 15) is the first metric, and balances directional and magnitude similarity between tensors and is a good general indicator of the similarity between two tensors. Linear-shape weighted V1 similarity (CLV1, Equation 17) is the second metric, and is defined as the inner product of the first eigenvector of the diffusion tensor (V1) from the template with V1 from the warped subject, weighted by the coefficient of linear shape (CL) for the template tensor (*i.e*., how similar V1 is, weighted by how informative V1 is). Planar-shape weighted V3 (CPV3, Equation 19) is the third metric, and is defined as the inner product of the third eigenvector of the diffusion tensor (V3) from the template with V3 from the warped subject weighted by the coefficient of planar shape (CP) for the template tensor (*i.e*., how similar V3 is, weighted by how informative V3 is). See Figure 1 for a visual depiction of CL and CP maps of the OMM-1 template. In areas where CL is large, the tensor shape is largely prolate (cigar-shaped), and therefore the direction of maximum diffusivity (*i.e*., V1) is well defined. In areas where CP is large, the tensor shape is largely oblate (plate-shaped), and therefore the direction of minimum diffusivity (*i.e*., V3) is well defined. CLV1 and CPV3 specifically probe how well orientational information has been aligned, which is most relevant in white matter regions. We expect those tools that consider rotational information (*i.e*., DTI data) during registration to perform best in these metrics.

**Figure 1:**
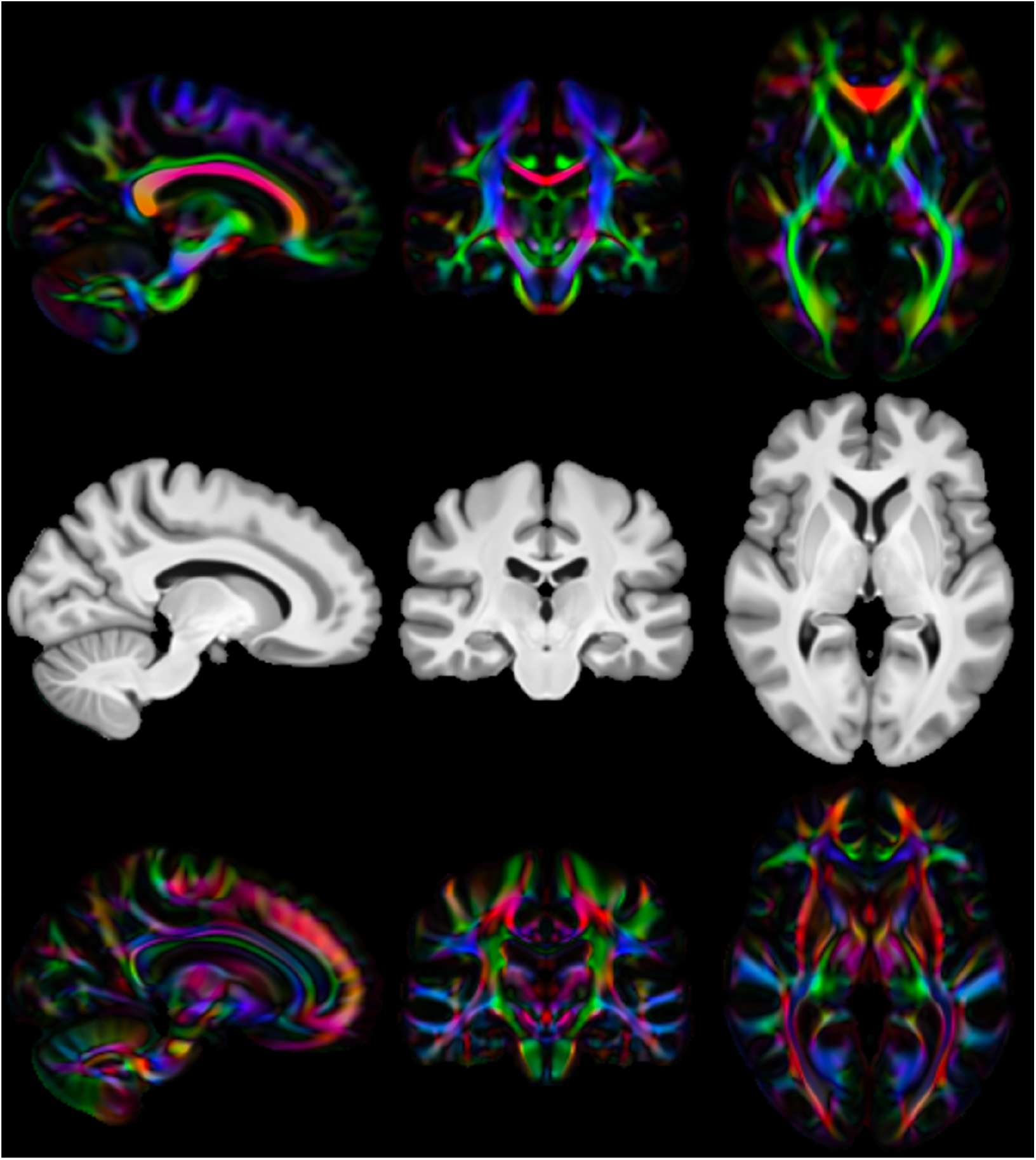
Visualisation of linear and planar shape coefficients in OMM-1 space: From top to bottom: Linear coefficient (CL) map of the template, T1w template for reference, planar coefficient (CP) map of the template. Images are directionally encoded colour maps of V1 and V3 respectively. Green = Anterior-Posterior, Red = Left-Right, Blue = Inferior-Superior. Both CL and CP values are highest in the white matter, but cover complementary regions therein. CLV1 is most sensitive to how well the primary diffusion direction is matched in voxels where diffusion occurs parallel to that axis only. CPV3 is most sensitive to how well the tertiary diffusion direction is matched in voxels where diffusion occurs within a plane defined by the primary and secondary diffusion directions. Together, these represent the voxels where tensor orientation can be reliably described by a single direction.

The fMRI accuracy metric used is task-fMRI cluster mass. This measures how consistently the registration tools are able to align those regions of the brain that are significantly activated (or deactivated) when performing a task. This assumes a general correspondence between brain structure and function, but this need not be exact. The benefit of this metric is that it is entirely independent of the modalities driving registration (*i.e*., T1w and DTI) and, therefore, there is little to no circularity in its interpretation, which cannot be said for the previous metrics.

Finally we calculate both size (| **J** |) and shape (CVAR—see Section 4.2.5 for definition) distortion metrics to evaluate how much each tool has had to deform the subject’s images to match the template. For any given level of accuracy (i.e., the preceding metrics) a smaller amount of distortion is usually preferred, as this indicates that the registration method is changing the original data as little as possible.

### 4.1. Ethics

All human imaging data used in this work are part of the open access Human Connectome Project Young Adult (HCP-YA) dataset. The data are pseudonymised and identifiable visual features, such as the face and ears, are obscured. Written informed consent to share the data “broadly and openly” was obtained for all participants by the original HCP-YA researchers (Elam et al., 2021).

### 4.2. Methods

Each of the following steps was performed independently for each registration method.

#### 4.2.1. Registration

All registration was performed to the OMM-1 template. This template contains both T1w and DTI volumes with isotropic resolution of 1 mm. Since both modalities were jointly aligned during template creation, they are intrinsically spatially consistent at the voxel level. It is therefore a good choice for registration with both unimodal and multimodal tools. Each tool was used to register all of the 100 unrelated HCP subjects to the template. FLIRT, FNIRT and ANTs used the T1w image only, while MMORF and DR-TAMAS used both the T1w and DTI images. The FLIRT affine matrix was used to initialise both FNIRT and MMORF, whereas ANTs and DR-TAMAS used their own affine registration methods. FNIRT and MMORF were run with custom configurations identified to empirically produce good results. ANTs and DR-TAMAS were run with slightly modified configurations that were found to improve registration accuracy over the defaults. All methods were run using a multi-resolution pyramid approach to a final warp resolution of 1 mm isotropic.

#### 4.2.2. T1w FreeSurfer Label Overlap

Automatically segmented subcortical (ASEG atlas, (Fischl et al., 2002, 2004)) and cortical (Destrieux 2009 atlas, (Destrieux et al., 2010)) parcellations for each subject were warped into template space. Jaccard coefficients (measuring label overlap) and Hausdorff distances (measuring the maximum error in label boundaries) of the corresponding warped parcellations were calculated for every possible pair of subjects. A pairwise approach was used because there are no target labels in template space. The average coefficient for each parcellation was then calculated across pairings for each subject, resulting in 100 values for each parcellation.

#### 4.2.3. DTI Similarity

Combined affine and nonlinear (apart from FLIRT) warp fields were used to resample each subject’s DTI volume into template space. Resampling of tensors was performed using the FSL tool vecreg, which includes preservation of principle direction (Alexander et al., 2001) reorientation, which is the most accurate way of accounting for the rotational effect of the warps when resampling DTI data. Tensors were then decomposed into three Eigenvalue- Eigenvector pairs (L1/2/3 and V1/2/3 respectively). OVL, CLV1 and CPV3 were then calculated between each subject’s warped DTI volume and the template. A pairwise approach was not necessary here since the DTI template volume itself acts as the target. CL and CP weighting coefficients (Alexander et al., 2000) were generated from the template DTI volume only. V1 and V3 similarity was calculated as the magnitude of the dot product of template and warped subject vectors in each template voxel. Similarity metrics were calculated per voxel according to the following formulae:

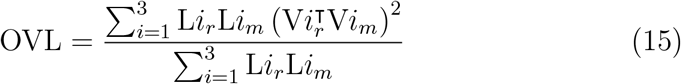

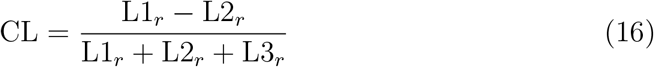

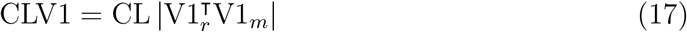

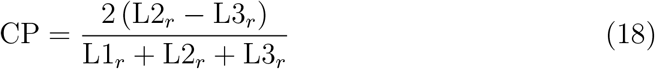

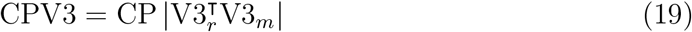

where the subscripts *r* and *m* represent the reference and resampled moving image respectively. All metrics were calculated within the template brain mask only.

#### 4.2.4. tfMRI Cluster Mass

The HCP task battery (Barch et al., 2013) consists of 7 tasks from which 86 contrasts are derived, and used to generate contrast of parameter estimate (COPE) images in subject T1w space. For each subject, the 86 COPE images were resampled into template space. FSL Randomise (Winkler et al., 2014) was then used to perform a group-level, non-parametric, ordinary-leastsquares, random-effects, one-group *t*-test on the mean COPE image (across subjects) for each contrast. The results of the group-level processing are 86 *t* -statistic maps/images and 86 family-wise-error (FWE) corrected *p*-value maps/images (one per contrast). The *t* -statistic map represents the group activation for a particular contrast, where more accurately aligned activations lead to higher *t* -statistics. The *p*-value map represents the statistical significance of the *t* -statistic at each voxel.

Cluster mass was then calculated as follows. Thresholding was applied to the FWE-corrected *p*-value map (at *p <* 0.05) for each contrast, to give a binary mask of significantly activated regions. This mask was then applied voxel-wise to the *t* -statistic map. The masked *t* -statistic map was then multiplied voxel-wise with a grey matter mask of the template, and summed across all voxels. The summed value is what we refer to as cluster mass, and is a single number per contrast and registration method. Cluster mass is, therefore, increased by both higher *t* -statistics and better alignment of grey matter. It can be calculated independently for each registration method, simplifying between-method comparisons

#### 4.2.5. Distortion

Two measures of distortion are considered, both evaluated within the template brain mask.

The first is the 5^th^ to 95^th^ percentile range of the log-Jacobian determinant (log | **J** |). This is a measure of volumetric distortion. Since the histogram of log | **J** | tends to be centred around zero, the mean is uninformative and, therefore, a range is more useful. This a fairly robust measure of the extent to which the local effect of the warp field is to expand/contract voxels (and is equally sensitive to both).

The second is the average cube-volume aspect ratio (CVAR, (Smith and Wormald, 1998)), defined as:

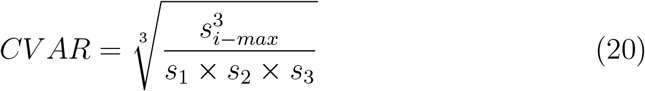

where *s*_*i*_ are the singular values of the local Jacobian matrix. CVAR represents the cube-root of the ratio of the volume of the smallest regular cube which can fully enclose the cuboid, to the cuboid’s own volume. Alternatively, for a deformation one can equally well define it as the cube-root of the ratio of the largest Jacobian singular value cubed, to the Jacobian determinant. This is a pure shape-distortion measure that is invariant to volumetric changes. For a perfect cube its value is ≥ 1, and it is greater than 1 for any shape where one or more sides of the cube are a different length to the others. Since it is always ≥ 1, the mean across voxels is a meaningful measure of the extent to which the local effect of the warp field is to alter the original shape of the underlying voxels.

Between them, these two measures present a good summary of the extent to which the warp is distorting the anatomy of the brain to achieve a given degree of registration accuracy. In general, lower distortion is for a given level of accuracy is preferred. When both distortion and accuracy increase then a more nuanced interpretation is required based on the relative changes in each.

### 4.3. Results

The results across all domains and metrics are summarised in Table 1. More detailed descriptions with accompanying figures are presented in the subsections that follow. Many figures utilise raincloud plots (Allen et al., 2021) that simultaneously show the raw data, a box and whiskers plot, and a density estimate. In all cases the interpretation of the box and whisker part of the plots is the same—the box shows the quartiles (25^th^, 50^th^ and 75^th^ percentiles), the whiskers extend to the final non-outlier datapoint, and the diamonds show outliers (any point more than 1.5 × the interquartile range from the edges of the box).

**Table 1:**
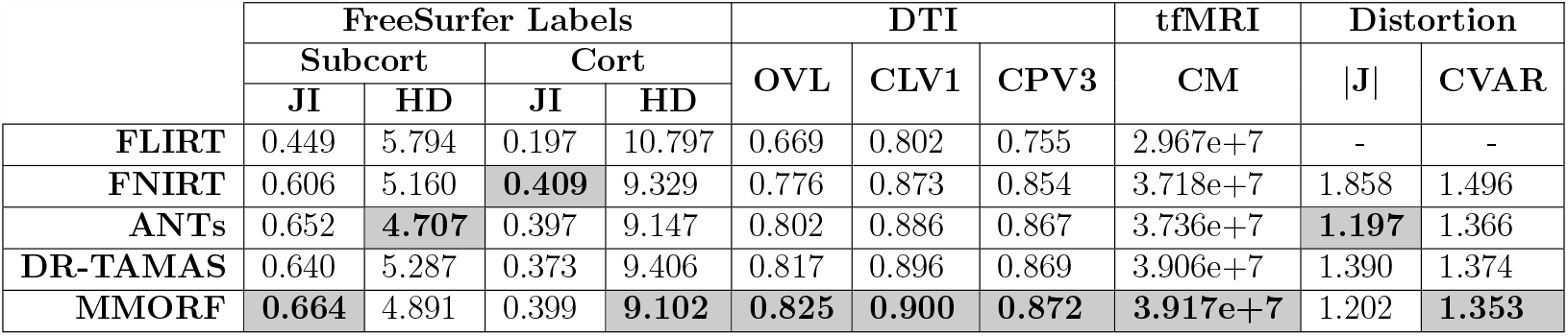
Summary of results for all domains. Structural (FreeSurfer labels): median Jaccard index (JI) and Hausdorff distance (HD) across subjects for both subcortical and cortical labels. Diffusion (DTI): median overal tensor similarity (OVL), linear coefficient-weighted V1 similarity (CLV1) and planar coefficient-weighted V3 (CPV3) similarity. Functional (tfMRI): Total cluster mass (CM) for all contrasts. Distortion: median 5^th^ to 95^th^ log-Jacobian determinant range (|**J**|) and cube-volume aspect ratio (CVAR) across subjects. The best performing method in each metric is highlighted.

#### 4.3.1. T1w FreeSurfer Labels

The Jaccard index and Hausdorff distance results are presented in Figures 2 and 3 respectively. In all cases, nonlinear registration leads to a major improvement in both metrics. MMORF and ANTs produce the best Jaccard index results in the subcortex, with FNIRT narrowly outperforming them in cortical regions. Hausdorff distance performance is slightly better in both subcortical and cortical regions for MMORF and ANTs. It is worth noting that subcortical contrast is relatively poor in T1w images from the HCP dataset, and certain structures (the left pallidum in particular) are poorly segmented in a number of subjects. This is the cause of the relatively heavy tails towards low Jaccard coefficients in Figure 2a.

**Figure 2:**
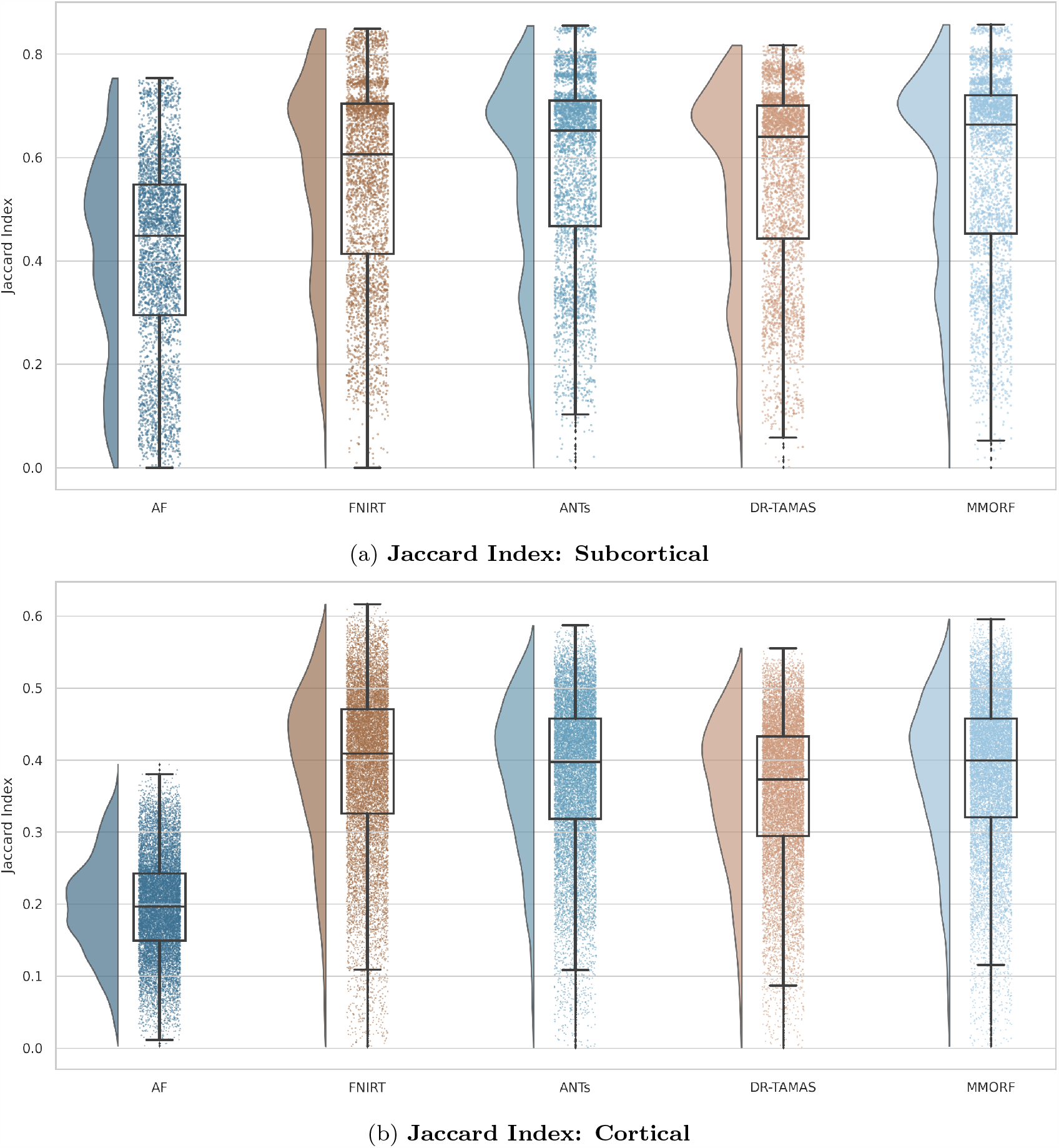
Subcortical (a) and cortical (b) Jaccard indices for FreeSurfer segmentation overlaps: All nonlinear methods improve over affine only, with the greatest improvement being in the cortex. Across all labels, MMORF and ANTs perform similarly (and best), with only FNIRT slightly outperforming them in the cortex.

**Figure 3:**
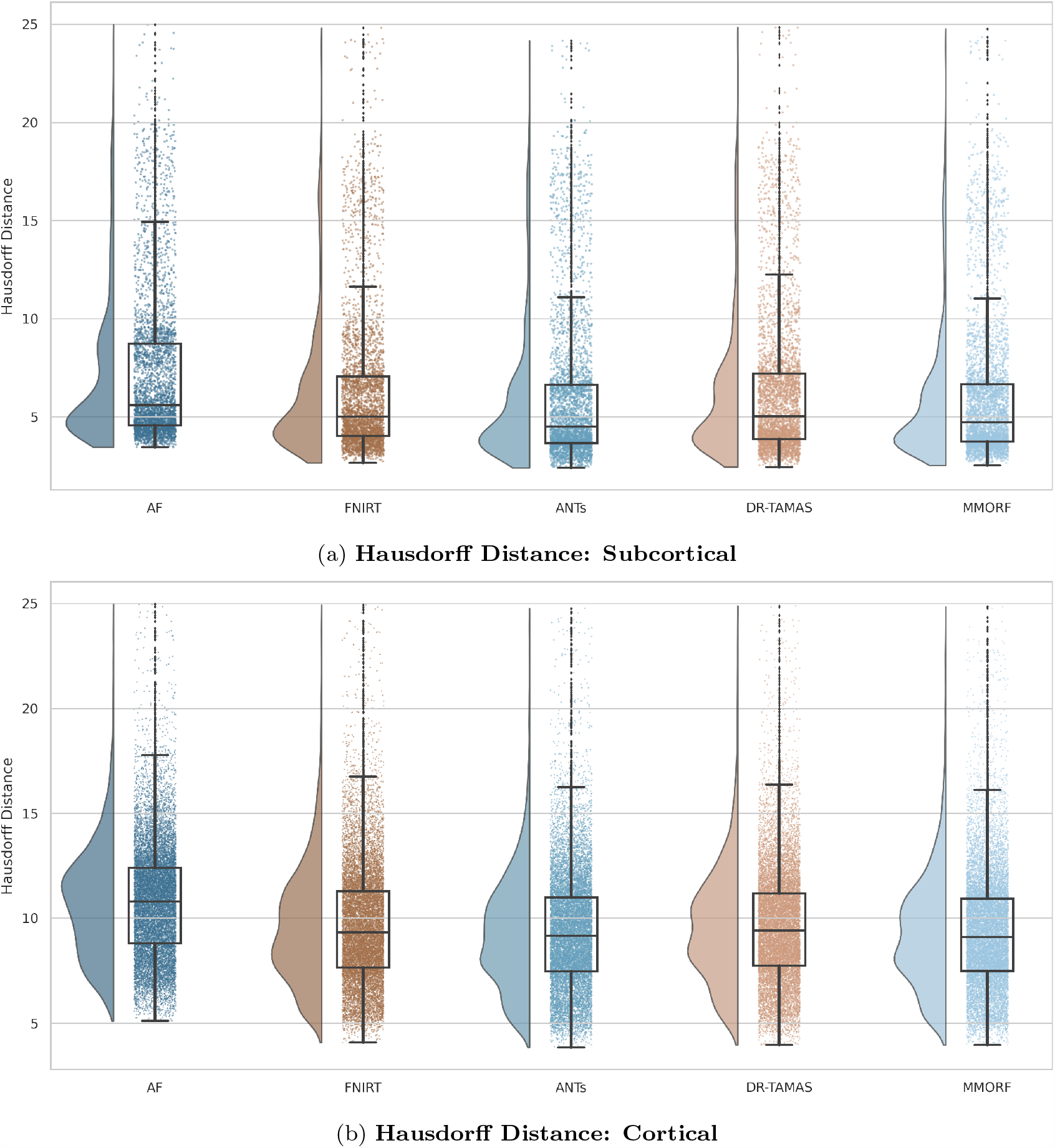
Subcortical (a) and cortical (b) Hausdorff distances for FreeSurfer segmentation overlaps: All nonlinear methods improve over affine only, with larger improvements evident in the cortex. Performance is very similar across all methods, with ANTs and MMORF slightly outperforming the others.

#### 4.3.2. DTI Similarity

OVL, CLV1 and CPV3 similarity results are presented in Figures 4 to 6 respectively. In all cases, the trend is the same: affine registration with FLIRT performs worst followed by FNIRT, ANTs, DR-TAMAS, and MMORF (in that order).

**Figure 4:**
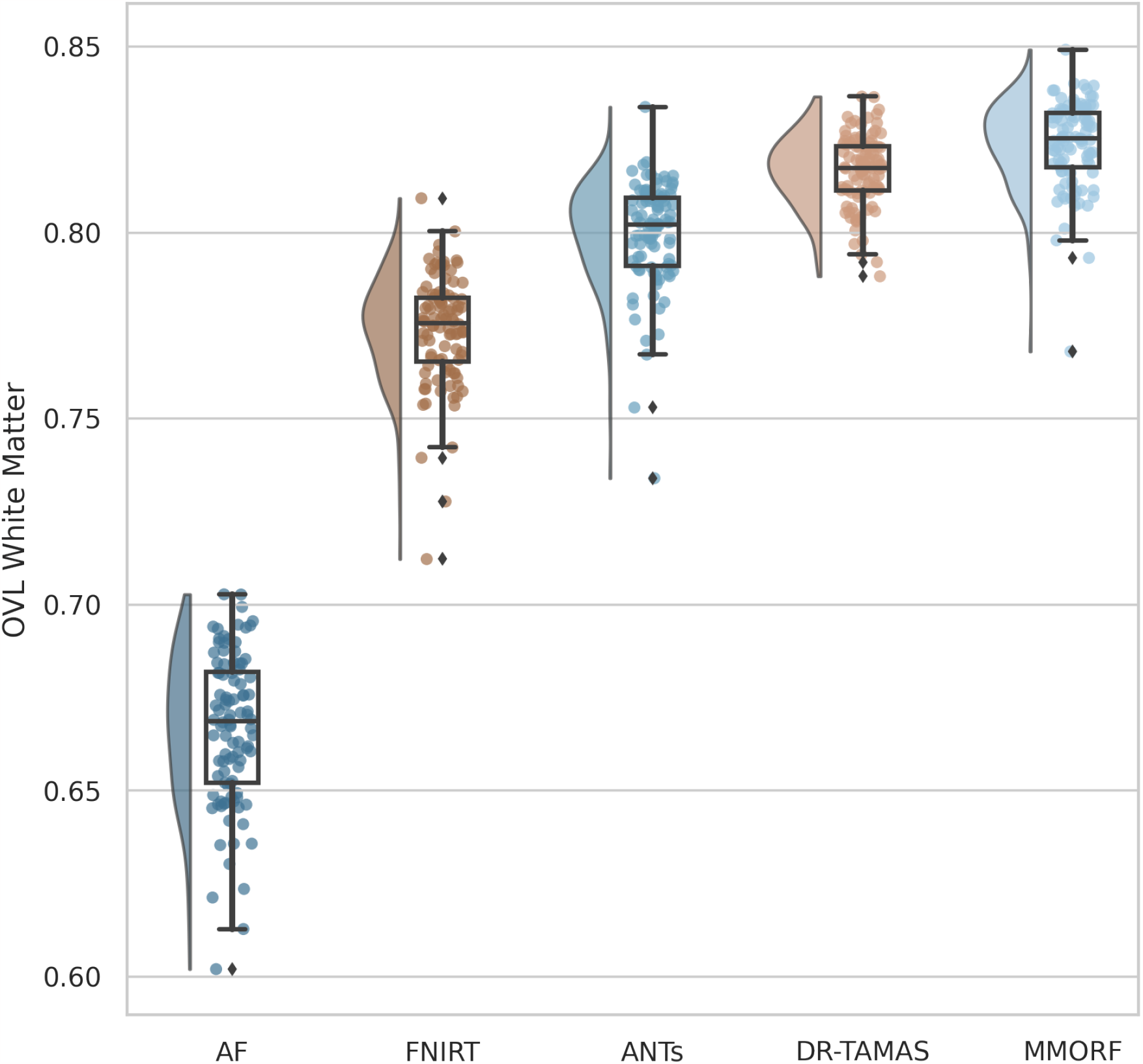
Overall tensor similarity (OVL): Calculated within a mask of the template white matter, defined as FA *>* 0.2. All nonlinear methods improve over affine only, with those methods that include DTI data in the registration (MMORF and DR-TAMAS) outperforming the T1w-only methods (FNIRT and ANTs). MMORF performs best overall.

**Figure 5:**
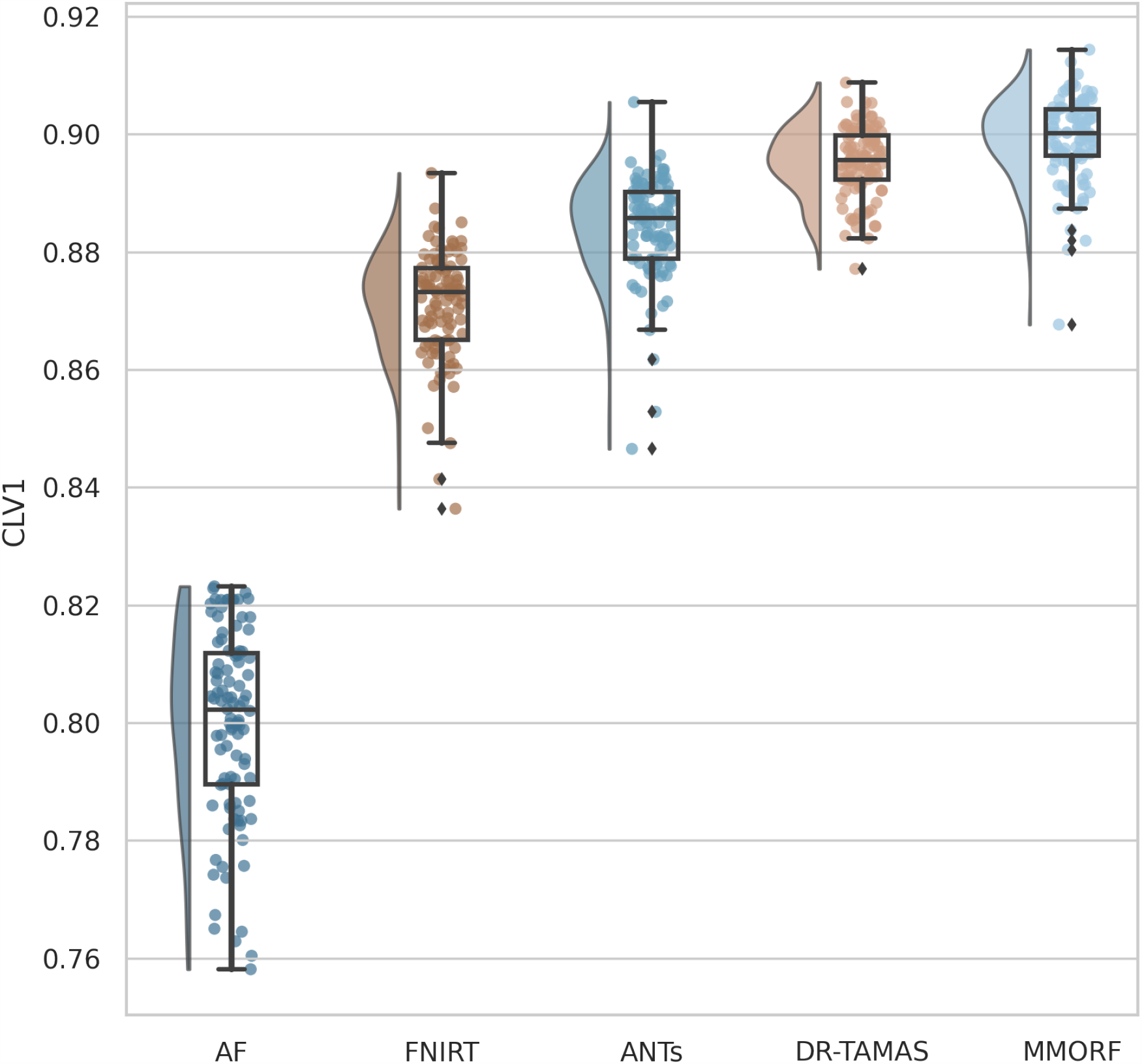
Linear shape weighted V1 similarity (CLV1): All nonlinear methods improve over affine only, with those methods that include DTI data in the registration (MMORF and DR-TAMAS) outperforming the T1w-only methods (FNIRT and ANTs). MMORF performs best overall.

**Figure 6:**
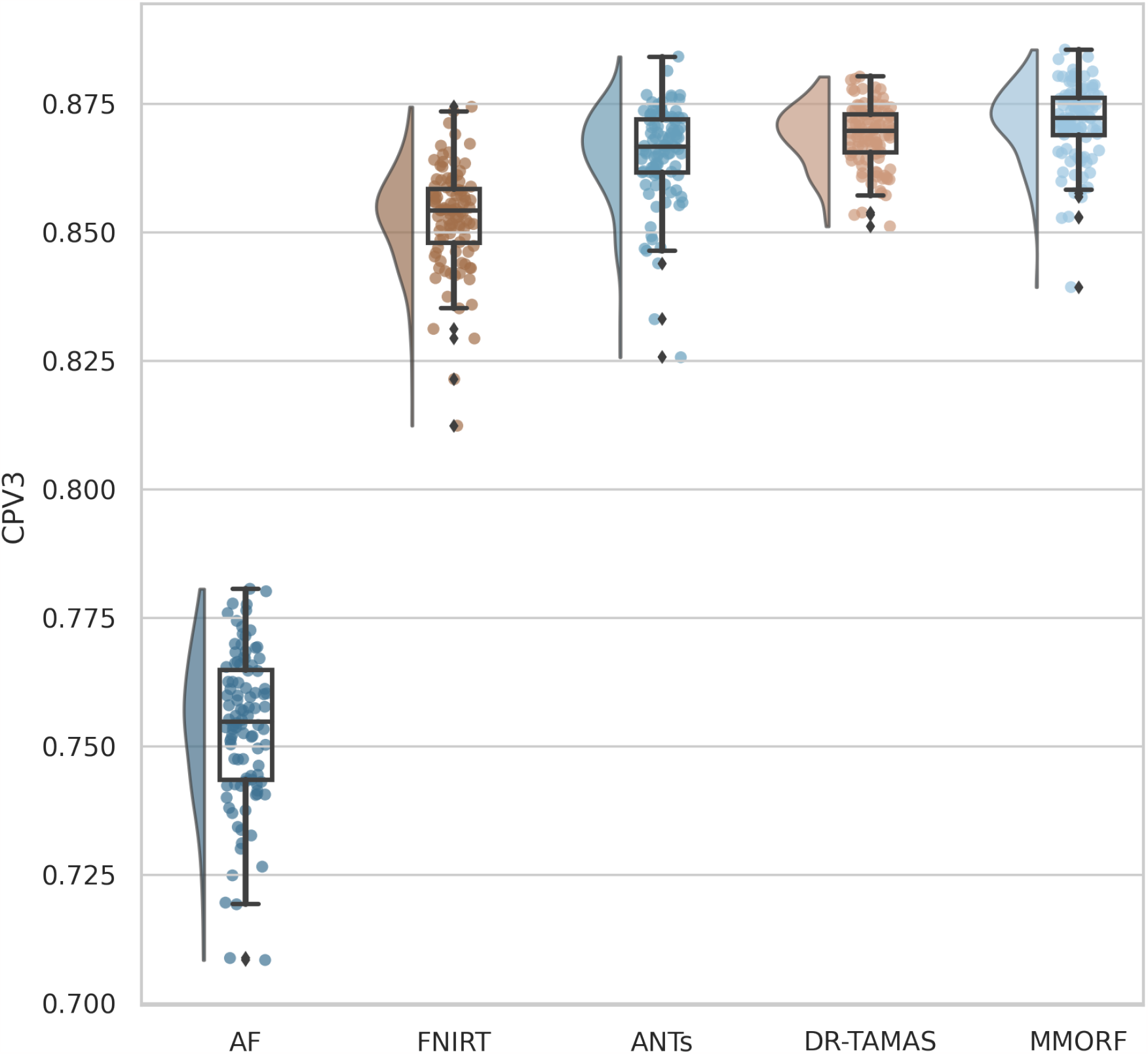
Planar shape weighted V3 similarity (CPV3): All nonlinear methods improve over affine only, with those methods that include DTI data in the registration (MMORF and DR-TAMAS) outperforming the T1w-only methods (FNIRT and ANTs). MMORF performs best overall.

#### 4.3.3. tfMRI Cluster Mass

Cluster mass results are presented in the form of percentage difference plots, and warrant some guidance on their interpretation. Each point on the plot represents one contrast. When comparing method *A* to method *B*, the x-axis represents the cluster mass of method *A*. The *x*-axis is log-transformed to account for the large range in cluster mass across contrasts. The *y*-axis represents the percentage improvement in cluster mass by method *A* over method *B*. Therefore, a point at location (*x, y*) = (10, 10) represents a contrast with a cluster mass of *exp*(10) for method *A*, and a 10 % improvement in cluster mass when using method *A* over method *B*. Similarly, a point at location (*x, y*) = (15, − 5) represents a contrast with a cluster mass of *exp*(15) for method *A*, and a 5 % reduction in cluster mass when using method *A* over method *B*. By choosing percentage difference for the *y*-axis we remove the bias towards large cluster mass contrasts, which would otherwise dominate if *y* were instead the simple difference in cluster mass between methods.

Cluster mass comparisons for MMORF vs FLIRT, FNIRT, ANTs and DR-TAMAS are presented in Figures 7 to 10 respectively. MMORF produces a large improvement across all contrasts compared to FLIRT, confirming that the CM metric is sensitive to improved registration accuracy. MMORF improves CM across most contrasts compared to FNIRT and, to a slightly lesser extent, ANTs. MMORF and DR-TAMAS produce very similar results for contrasts with high CM (towards the right of the x-axis), but MMORF performs consistently better for small and medium CM contrasts.

#### 4.3.4. Distortion

Comparisons of the 5^th^ to 95^th^ percentile Jacobian determinant range (volumetric distortion) and the average CVAR (shape distortion), which are both evaluated only within a brain mask, are presented in Figures 11 and 12 respectively. FNIRT shows significantly more distortion than the other methods. ANTs and MMORF display similar levels of volumetric distortion, that are lower than DR-TAMAS. MMORF displays the least shape distortion, followed by ANTs and DR-TAMAS. The difference between ANTs and DR-TAMAS is likely due to larger deformations within the white matter with DR-TAMAS, due to diffusion information driving the registration harder in those regions.

**Figure 7:**
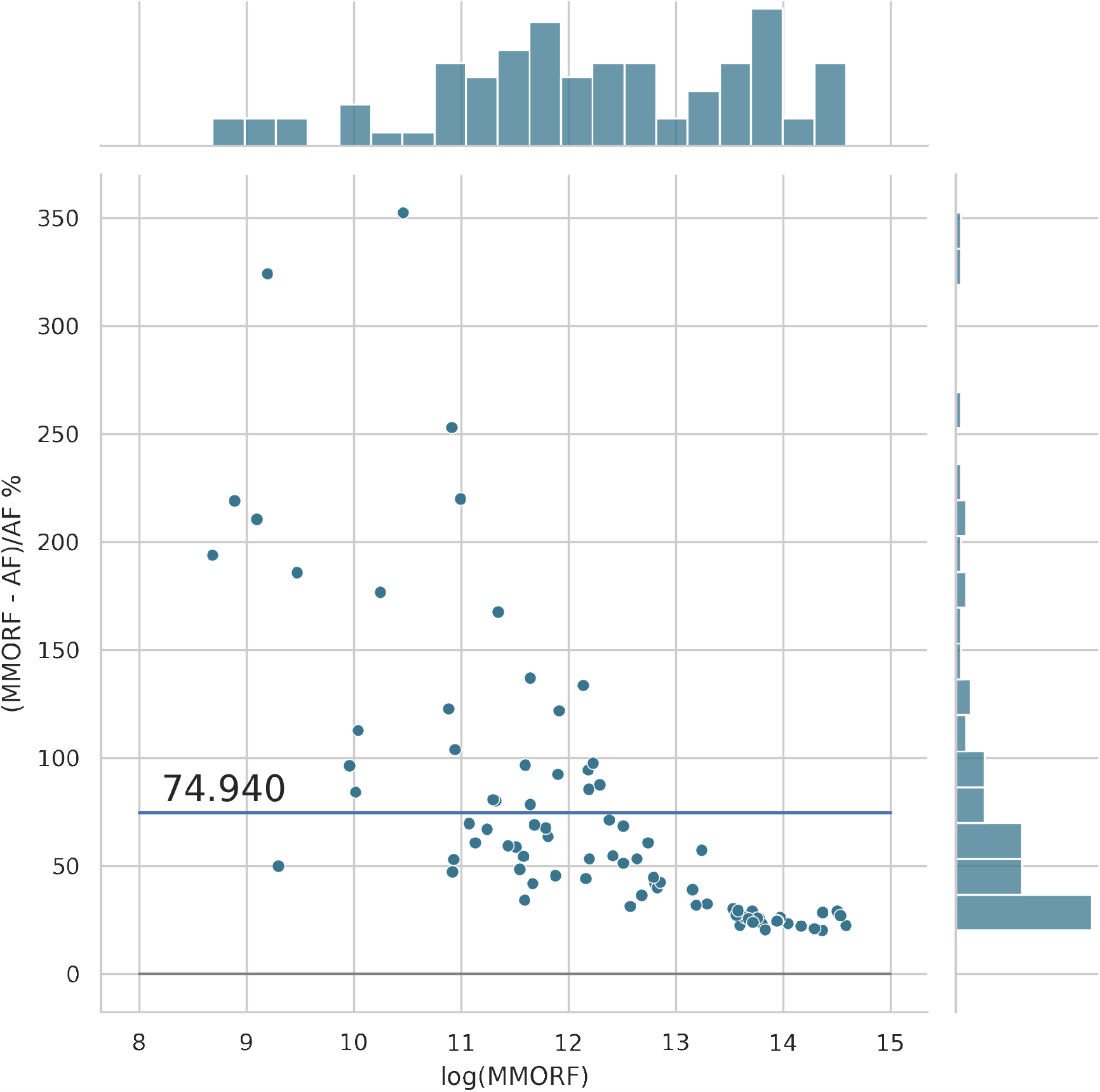
MMORF vs FLIRT cluster mass: Across clusters of all size, MMORF (nonlinear) outperforms FLIRT (linear), with an average improvement in cluster mass of ≈75%. This is not surprising, but serves to demonstrate that cluster mass is sensitive to registration accuracy.

**Figure 8:**
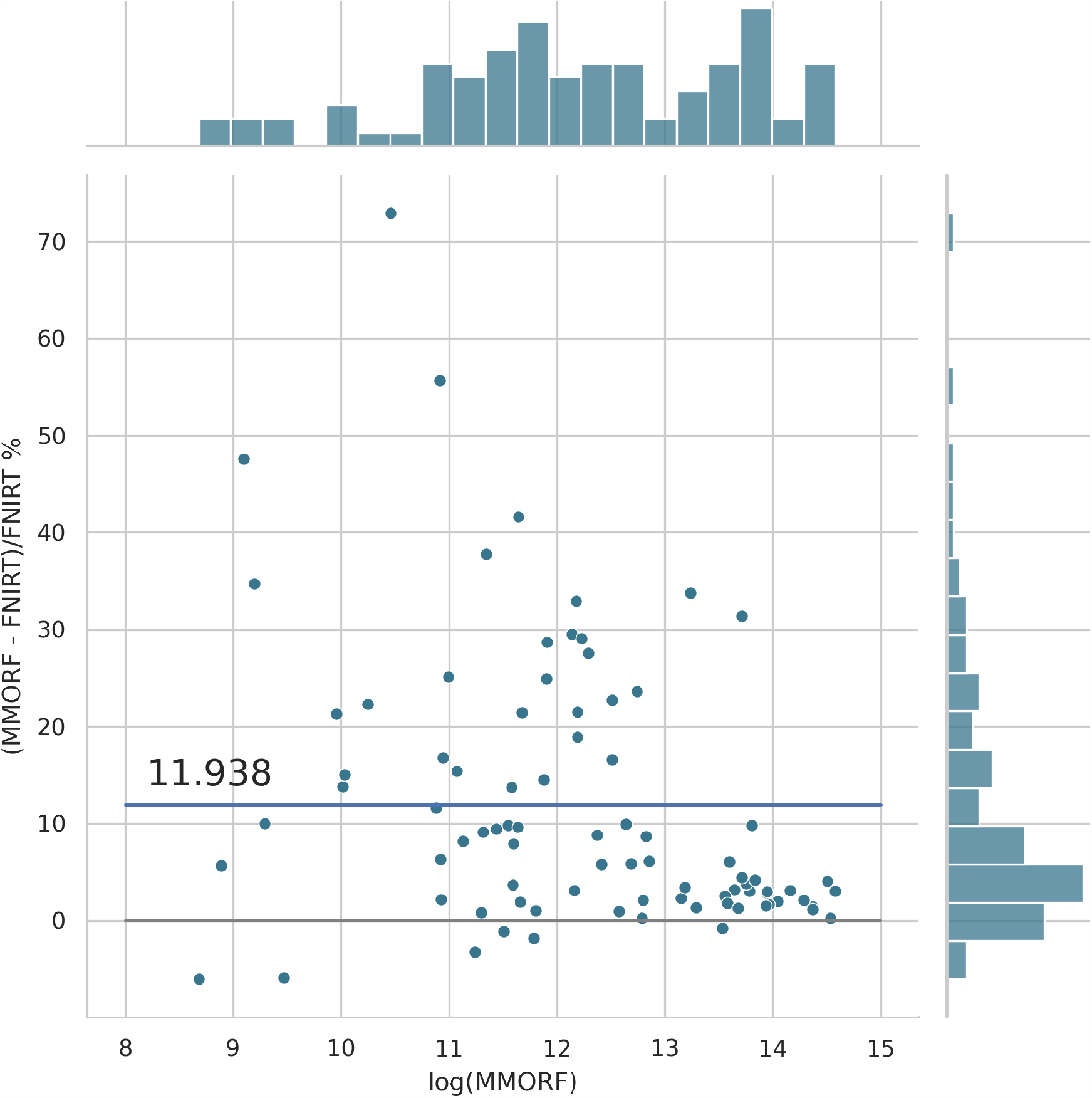
MMORF vs FNIRT cluster mass: MMORF outperforms FNIRT with an average improvement in cluster mass of ≈12%. The largest improvements are in the mid-sized clusters.

**Figure 9:**
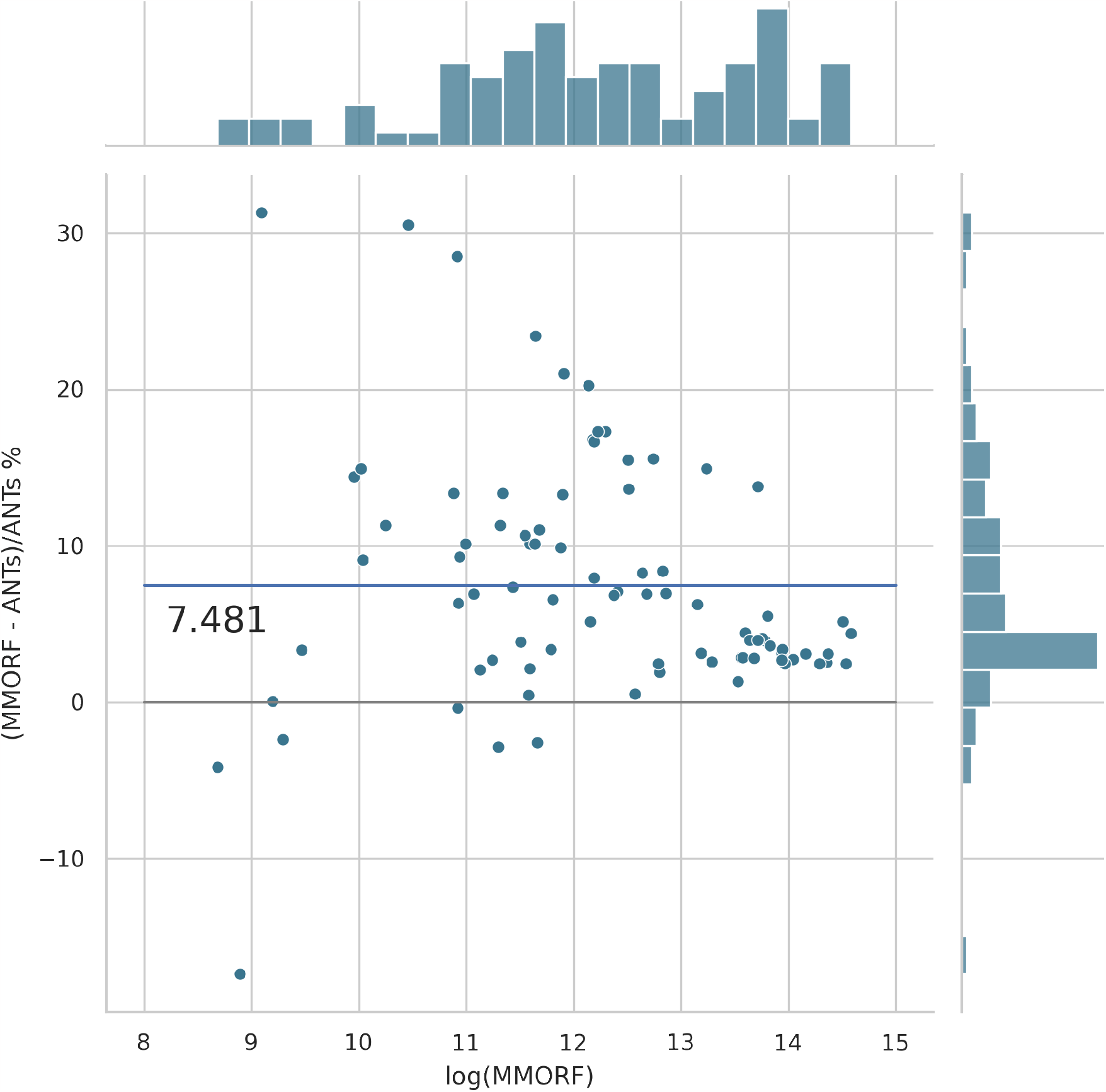
MMORF vs ANTs cluster mass: MMORF outperforms ANTs with an average improvement in cluster mass of ≈ 7.5%. The largest improvements are in the mid-sized clusters.

**Figure 10:**
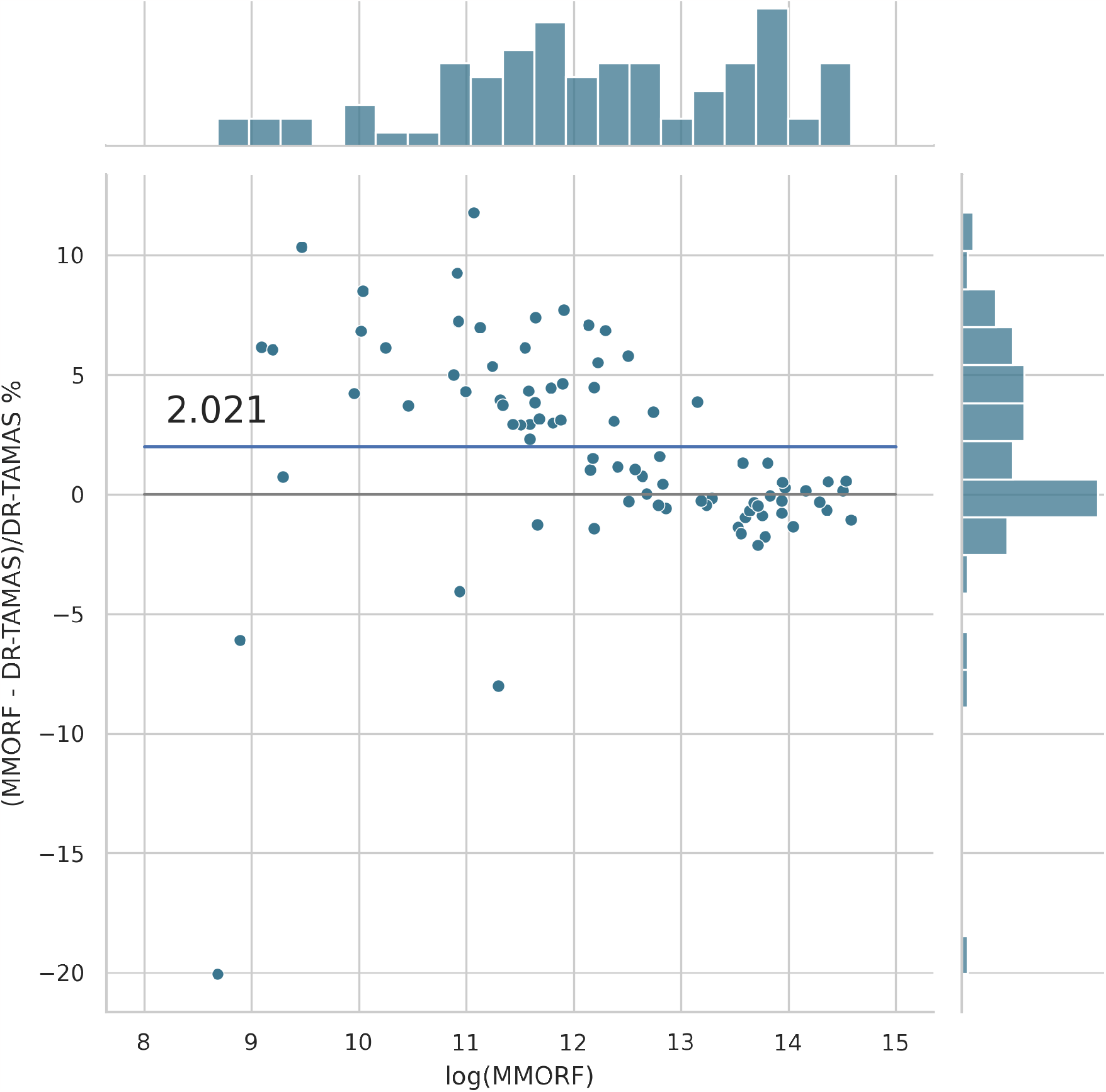
MMORF vs DR-TAMAS cluster mass: MMORF outperforms DRTAMAS with an average improvement in cluster mass of ≈2%. The largest improvements are in the mid-sized clusters.

**Figure 11:**
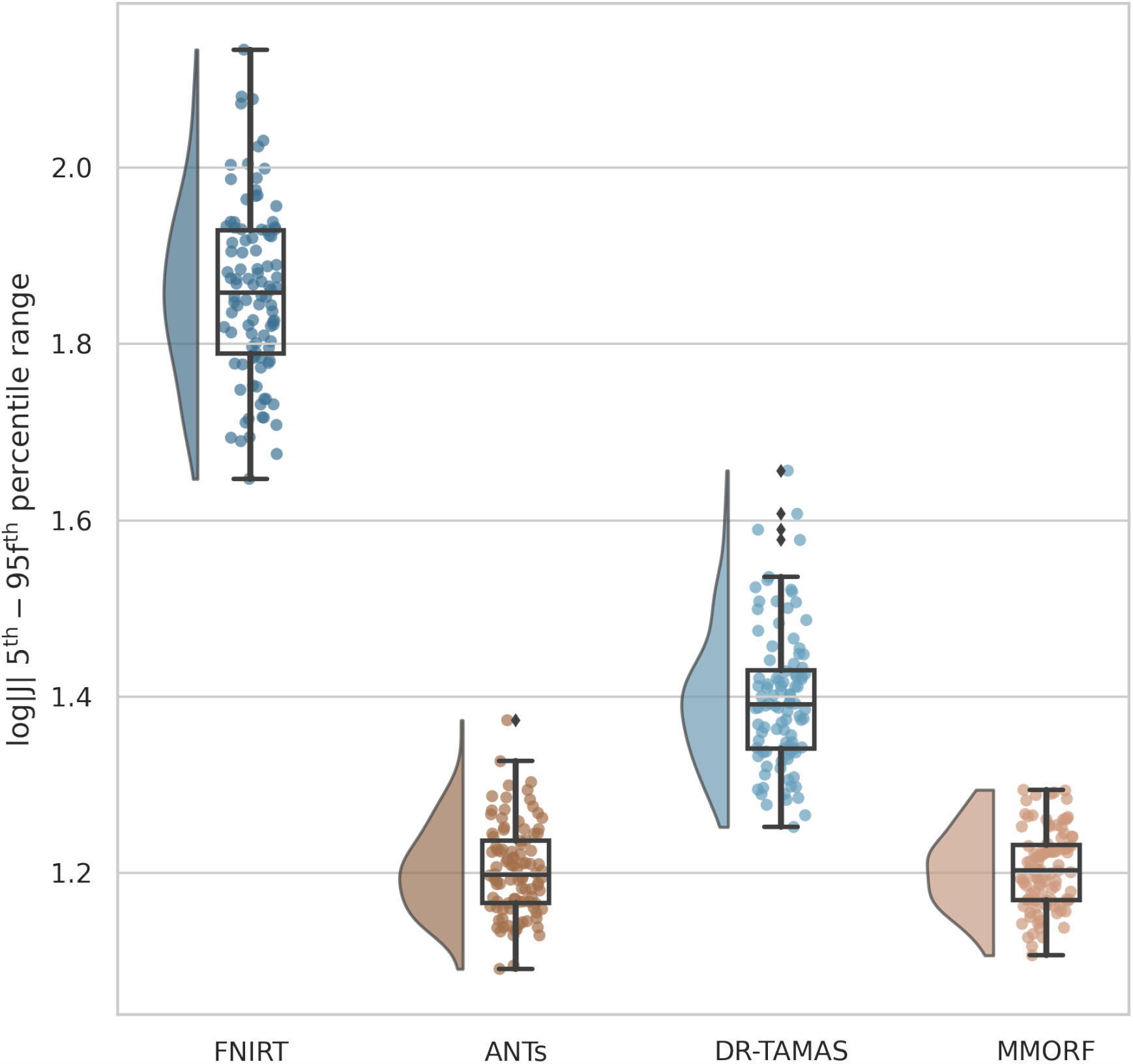
**5**^**th**^ **to 95**^**th**^ percentile Jacobian determinant range: MMORF and ANTs have the lowest, and very similar, levels of volumetric distortion. This is despite the fact that the MMORF warps are also trying to align the DTI information in the white matter, which would be expected to increase the amount of distortion (as can be seen has happened for DR-TAMAS). FNIRT shows the largest amount of distortion.

**Figure 12:**
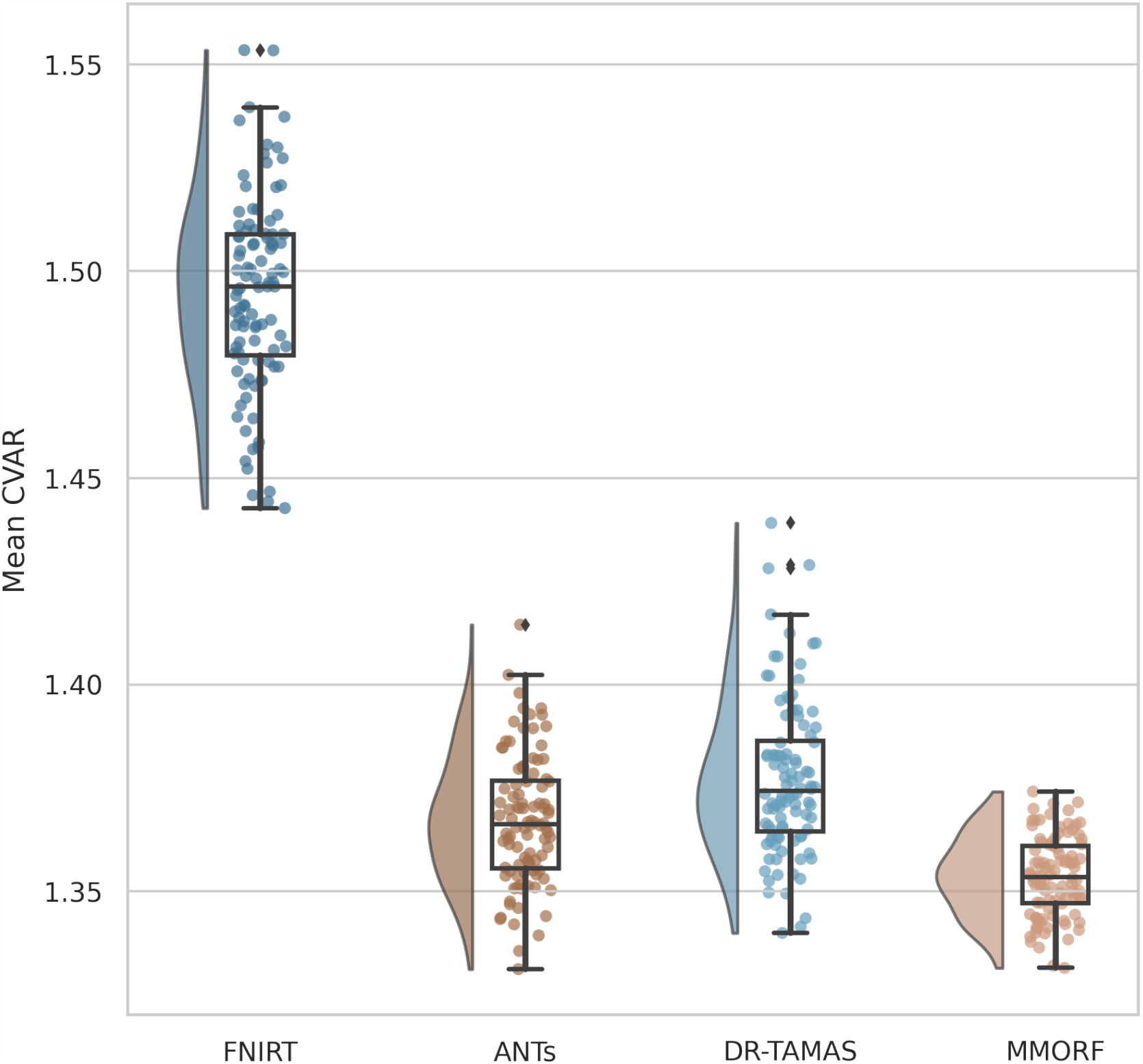
Mean cube-volume aspect ratio (CVAR) shape distortion: MMORF produces the lowest level of shape distortion on average, followed by ANTs, DR-TAMAS and FNIRT. As with the volumetric distortion, this is in spite of the fact that the MMORF warps are also trying to align the DTI information in the white matter, which would be expected to increase the amount of distortion (as can be seen has happened for DRTAMAS). This also demonstrates that MMORF’s low levels of volumetric distortion (see Figure 11) do not come at the cost of increased shape distortion, which can be observed when using a regularisation that penalises only the Jacobian determinant (Lange et al., 2020).

### 4.4. Summary

Of the methods tested, MMORF is the most consistently high-performing across the full range of evaluation metrics.

It might be expected that the T1w-only-driven registration methods would perform well when evaluated with a T1w-derived similarity metric (particularly in the cortex) and, indeed, both FNIRT and ANTs perform very well in the label overlap metric in this region. It is also possible that the inclusion of DTI information during registration might negatively affect such a metric and, again, we do indeed see that DR-TAMAS performs relatively poorly in the cortex, despite good subcortical performance. MMORF, however, does not seem to suffer in the same way—performing on par with ANTs both cortically and subcortically. We may, therefore, conclude that MMORF is a good choice of method, on par with ANTs, when a structurally-derived segmentation comparison is the type of study for which registration is being employed.

The value of including DTI information in the registration is clear, if unsurprising. MMORF and DR-TAMAS noticeably outperform both FNIRT and ANTs across all DTI similarity metrics. Of the multimodal registration methods, MMORF has the advantage over DR-TAMAS across all metrics. The OVL metric shows that, within the white matter, both the shape and the size of the tensors are better matched by MMORF. The CLV1 and CPV3 metrics show that the most informative directions of the tensor are also best aligned by MMORF.

We believe the tfMRI results to be the most unbiased assessment of registration accuracy presented here, since it is evaluated using a modality that was never seen by any of the registration methods being tested. Since MMORF outperforms all other methods under test, this is the clearest indication of its high registration accuracy. It is notable that FNIRT, the best performing method in terms of cortical label overlaps, does not perform as well in this (also cortical) metric. Similarly, DR-TAMAS is the closest performing method to MMORF in the tfMRI evaluation, but is the poorest performing method in cortical label overlap. This is a clear example of the benefit in not only considering the more circular evaluation metrics (i.e., those derived from the same modalities that are driving the registration) when comparing registration tools.

Finally, in terms of distortion, MMORF demonstrates levels that are similar to or lower than those of ANTs, despite matching it in label overlap and bettering it in both the DTI and tfMRI metrics. DR-TAMAS produces more distortion on average than its sibling method, ANTs, which is not unexpected given that DR-TAMAS is also being driven by DTI information in the white matter. FNIRT produces larger distortions than the other nonlinear methods. Without MMORF for comparison, this might easily be interpreted as the inevitable consequence of driving a small deformation framework method very hard to maximise accuracy metrics. However, given that MMORF and FNIRT have the same transformation model, this instead highlights the value of using a more biologically plausible regularisation model within the smalldeformation framework.

## 5. Discussion

The primary objective in developing MMORF was to exploit the rich information available in multimodal datasets in order to align brain images with maximal accuracy. A truly multimodal tool must be able to leverage information about both magnitude and directionality in each voxel, since we are able to capture and represent both of these with MRI. To this end, MMORF is able to explicitly, and simultaneously, optimise both the displacement and rotational effects of a warp field. Importantly, not *optimising* for rotation (as is the case for any scalar registration method) does not mean that there *is* no rotation, only that it cannot be controlled to either improve directional alignment or prevent the introduction of misalignment. Simultaneously optimising over multiple modalities through a single warp both allows the unique information from each modality to influence the resulting transformation (as when each modality is aligned separately), and ensures that all modalities remain co-registered following warping (as when using only a single modality to drive alignment).

But even when combining information across modalities, image registration is still a highly underdetermined problem and, therefore, alignment accuracy (as measured by how well anatomy are co-localised) is highly dependent on the choice of warp regularisation. We therefore included in MMORF a regularisation model that promotes highly biologically plausible deformations, thereby effectively controlling excessive levels of distortion in both shape and size.

The combination of our multimodal approach and our regularisation metric is computationally intensive, particularly when it comes to calculating the Hessian matrix required by MMORF’s Gauss-Newton optimisation strategy. We address this through GPU parallelisation of the most computationally intensive parts of the algorithm, allowing MMORF to run to 1 mm warp resolutions within 5 to 45 minutes (depending on image resolution and number of modalities).

We have evaluated MMORF across four domains—T1w-label overlap, DTI similarity, tfMRI cluster mass and image distortion. Performance was benchmarked against three established registration tools—FNIRT, ANTs, and DR-TAMAS. These tools represent MMORF’s predecessor, a stateof-the-art unimodal method, and the most similar multimodal alternative, respectively. A common theme was that methods performed well when tested on metrics derived from modalities used in their respective cost functions—e.g., FNIRT performed well on T1w-label overlap, and DR-TAMAS performed well on DTI similarity. MMORF performed best or near-best in both scalar and tensor evaluations, best in the held-out tfMRI evaluation, and best in terms of distortion.

FNIRT was able to slightly outperform MMORF in cortical T1w-label overlap, but was outperformed by MMORF across all other metrics. FNIRT induces the most distortion out of the methods tested. This likely contributed to the relatively poor DTI similarity results, since excessive deformations are likely to cause incorrect rotation of the tensors. Interestingly, having the best cortical label overlap has not translated to the best tfMRI performance, with FNIRT actually showing the poorest performance, despite both domains being evaluated on the cortex. The strong cortical T1w-label performance is, therefore, likely due to overfitting of the T1w image similarity metric, and highlights the importance of holistically evaluating registration performance, including the use of a held-out modality. These results demonstrate the significant improvement in performance of MMORF over its predecessor. Since both the inputs and outputs to MMORF are fully compatible with FNIRT and FSL, the benefits of this improvement can be realised by simply substituting in the new method.

ANTs performed near-identically to MMORF in both cortical and subcortical T1w-label overlap measures, making them the best performing methods in this domain. The inclusion of DTI data in MMORF registration has, therefore, not compromised the alignment of the T1w channel, which is not necessarily a given (see the comparison to DR-TAMAS below). The benefits of including the DTI data are evident in the DTI similarity results, where MMORF clearly has the advantage over ANTs. MMORF consistently outperforms ANTs in the tfMRI evaluation, indicating better anatomical consistency in grey matter, despite similar label overlap performance. Levels of distortion are very comparable between these two methods, despite there being more information to drive white matter deformations harder in MMORF.

DR-TAMAS was the poorest performing nonlinear method in terms of cortical T1w-label overlap, and second-poorest subcortically. Since it largely shares its scalar registration algorithm with ANTs, this suggests that this is due to the inclusion of DTI data in driving the registration. As has already been noted, this was not the case for MMORF. This could therefore be attributed to the differences in DTI cost functions employed by DR-TAMAS and MMORF—where MMORF uses the whole tensor, while DR-TAMAS uses a combination of the mean diffusivity (a scalar) and the deviatoric tensor (shape and direction information only). DR-TAMAS came a close second to MMORF across all DTI similarity metrics, with performance clearly better than the scalar-only methods. DR-TAMAS was also the closest performing method to MMORF in the tfMRI evaluation, demonstrating again that cortical segmentation performance does not necessarily translate to well matched cortical activations. In terms of amount of deformation, DR-TAMAS induced more deformations than MMORF, despite having been outperformed across all accuracy measures.

One limitation of this and, in fact, most evaluations of registration methods is that we are likely to be able to better tune our own method than those against which we are comparing. However, starting from the default settings, we have made an effort to optimise the performance of each, and kept any adjustments that proved beneficial. As such, we believe that our results are representative of what can be expected in real-world use. While there was always at least one tool with similar performance to MMORF in each test, none were consistently as high-performing across the board.

Note that, although our primary objective was human brain alignment, MMORF does not rely on any human brain priors (*e.g*., tissue maps or assumed brain size), and may be applied to any domain of medical imaging. For example, MMORF has been successfully used in the generation of multimodal, non-human primate brain templates of several phylogenetically distant species for the purpose of performing comparative anatomy of white matter tracts (Roumazeilles et al., 2022). Similarly, while the HCP enables us to compare the relative performance of each registration tools across multiple imaging modalities using a single dataset, our evaluation here is largely confined to the cerebrum as there is little to no labelled data in the cerebellum, brain stem, and spinal cord. MMORF may, in fact, be particularly beneficial for studying these non-cerebral regions, and we hope to explore this is in future.

In this work we have only considered MMORF’s performance in comparison to other volumetric registration tools. There is strong evidence that using surface-based registration has a beneficial effect on cortical alignment accuracy (Coalson et al., 2018), and we are not advocating the use of MMORF instead of those methods when performing cortical or surface-based cross-subject fMRI analyses, for example. However, volumetric registration is, as yet, still the only way to align non-cortical brain regions, and is likely to remain just as valuable as surface registration in neuroimaging research for the foreseeable future, especially when data are registered intra-individually and when surface reconstructions fail, for example, due to the presence of intra-axial lesions (such as brain tumours).

Given MMORF’s performance, reasonable execution times, and simple compatibility with FSL, we believe it to be an excellent choice of tool for volumetric registration—regardless of the domain of any follow-on analysis. As such, it is easy to recommend MMORF to all users working with multimodal neuroimaging data, and in particular those who already rely on FSL for their analyses. They will benefit from state-of-the-art registration accuracy across all domains of their downstream analyses, with very little modification to any existing pipelines. MMORF is available as part of FSL in releases newer than 6.0.7.

## 6. Data and Code Availability

MMORF binaries and the associated source code are available for download via the standard FSL installer3. A stand-alone Singularity image, instructions for running MMORF, and example configuration files are available on the FMRIB GitLab server4. All analysis scripts, figures, figure source code and latex source code that were used to generate this manuscript are available on the FMRIB GitLab server5.

## 7. Author Contributions

**Frederik Lange:** Conceptualisation, Methodology, Software, Validation, Formal analysis, Investigation, Writing (original draft, review and editing), Visualisation. **Christoph Arthofer:** Methodology, Software, Validation, Writing (review and editing). **Andreas Bartsch and Gwenaëlle Douaud:** Validation, Writing (review and editing). **Paul McCarthy:** Software, Data Curation. **Stephen Smith:** Conceptualisation, Methodology, Validation, Resources, Writing (review and editing), Supervision, Project administration, Funding acquisition. **Jesper Andersson:** Conceptualisation, Methodology, Software, Validation, Formal analysis, Writing (review and editing), Supervision, Project administration.

## 8. Acknowledgements

This research was funded by a Wellcome Trust Collaborative Award (215573/Z/19/Z). The Wellcome Centre for Integrative Neuroimaging is supported by core funding from the Wellcome Trust (203139/Z/16/Z). Computation used the Oxford Biomedical Research Computing (BMRC) facility, a joint development between the Wellcome Centre for Human Genetics and the Big Data Institute supported by Health Data Research UK and the NIHR Oxford Biomedical Research Centre. The views expressed are those of the author(s) and not necessarily those of the NHS, the NIHR or the Department of Health. Data were provided in part by the Human Connectome Project, WU-Minn Consortium (Principal Investigators: David Van Essen and Kamil Ugurbil; 1U54MH091657) funded by the 16 NIH Institutes and Centers that support the NIH Blueprint for Neuroscience Research; and by the McDonnell Center for Systems Neuroscience at Washington University.

## 9. Declaration of Competing Interests

We have no competing interests to declare.

## Footnotes

1 https://fsl.fmrib.ox.ac.uk/fsl/fslwiki/FDT/UserGuide#DTIFIT

2 DOI 10.17605/OSF.IO/S9GE4

3 https://fsl.fmrib.ox.ac.uk/fsl/fslwiki/FslInstallation

4 https://git.fmrib.ox.ac.uk/flange/mmorf_beta

5 https://git.fmrib.ox.ac.uk/flange/mmorf_toolbox

## Notes

### Competing Interest Statement

The authors have declared no competing interest.

